# Kv4, Kv2, and Kv3 currents shape intrinsic lateral olivocochlear excitability independent of hair cell dysfunction during development and ageing

**DOI:** 10.64898/2026.07.02.736013

**Authors:** Adam J. Carlton

## Abstract

Intrinsic lateral olivocochlear (iLOC) neurons provide vital brainstem efferent feedback to the cochlea in order to modulate hearing sensitivity through synapses onto type-I spiral ganglion neurons. During ageing or mutations affecting hair cell transduction in mice, efferent neurons rewire to form direct axo-somatic synapses onto inner hair cells (IHCs), recapitulating a synaptic configuration typically only restricted to the immature cochlea. Whether this rewiring reflects a compensatory mechanism or some form of attempted repair, or how iLOC biophysics change throughout ageing and this rewiring process, is not known. We utilised whole-cell patch-clamp electrophysiology to investigate iLOC activity and their underlying biophysics across the wild-type mouse lifespan. We show that iLOC neurons undergo a progressive increase in excitability with post-natal development and ageing, producing more spikes for a given stimulus. This intrinsic excitability shift was driven by the developmental decline in the A-type Kv4 mediated potassium current and increase in Kv2 mediated current. In ageing animals, and distinct from post-natal development, further increased firing rates were supported by an increased size of the fast-activating Kv3 current. Spontaneous bursting activity remained present in ageing iLOC neurons, and no reversion to an immature biophysics profile was evident. Interestingly, despite robustly eliciting efferent rewiring of IHCs, an accelerated ageing-like re-innervation genetic model did not recreate the biophysical changes in the iLOC neurons that reflected the ageing system. This work reveals distinct processes occurring within the iLOC feedback system, and shows that age-related enhancements of SGN resting activity are not triggered by deficits in IHC transduction.

**Significance Statement:** Modulation of cochlea output from the brainstem remains poorly understood, and the efferent feedback from the intrinsic lateral olivocochlear (iLOC) neurons has been shown to be vital for maintaining hearing sensitivity after noise exposure. Efferent neurons undergo significant rewiring within the cochlea during maturation that appears to revert to an immature state with ageing or with dysfunction of the inner hair cell sound-transducing apparatus, though how iLOC activity adapts to this is unknown. Here we show that ageing involves distinct shifts in sodium and potassium current biophysics that support the generation and development of spontaneous bursting activity. Disrupting sound transduction revealed that iLOC age-related biophysical changes and cochlear rewiring are independent, and defective transduction is insufficient to recreate ageing biophysics.

## Introduction

Inner hair cells (IHCs) are the primary sensory receptor of the cochlea, and are responsible for the transduction of all perceivable sound. This is reflected in their innervation, with each IHC being contacted by around 20 unbranched and myelinated type I spiral ganglion neurons (SGNs), which faithfully transmit sound information to the brain **(Keithley and Feldman, 1982; Burda and Branis, 1988; Pujol et al., 1998**). The firing rates of these SGNs are dictated by the balance between their intrinsic biophysical properties, spontaneous synaptic release from IHCs, and excitatory or inhibitory modulation from the central nervous system (**Darrow et al., 2006b**). Feedback to the cochlea from the brainstem is poorly understood, and by modulating cochlear output the olivocochlear efferent system is thought to assist with sound localisation, selective attention, and hearing in noise (**Groff and Liberman, 2003; Darrow et al., 2006a**). One primary source of this modulation is the intrinsic lateral olivocochlear (iLOC) efferent system, arising from the tonotopically organised lateral superior olive (LSO) within the brainstem. By default, iLOC neurons are primarily cholinergic and excitatory to SGNs (**Felix and Ehrenberger, 1992; Safieddine and Eybalin, 1992**), but upregulate inhibitory dopamine following loud sound exposure (**Wu et al., 2020**). Ablation of iLOC feedback results in a reduced capacity to recover hearing thresholds after exposure to loud noise, suggesting iLOC feedback is important for maintaining auditory sensitivity following exposure to damaging sound levels. **(Sitko et al., 2025**).

Alongside this remarkable neurotransmitter plasticity, iLOC neurons also show plasticity in terms of their synaptic targets. In the pre-hearing cochlea, prior to post-natal day (P) 12 in mice, cholinergic projections originating from the medial olivocochlear system directly contact IHCs with an inhibitory cholinergic synapse which regulates IHC spontaneous activity and properly patterns the downstream auditory pathway (**Clause et al., 2014; Wang et al., 2021**). After P12 these direct axo-somatic synapses onto the IHCs detach, and at the same time-point iLOC neurons acquire their mature configuration, synapsing onto SGNs selectively. Concurrently, through an unknown mechanism, as iLOC neurons mature they develop calcium-dependent spontaneous bursting activity, and the duration of these bursts can be increased by noise exposure (**Hong et al., 2022; Hong and Trussell, 2025**). This activity is hypothesized to provide continuous excitatory input to the SGNs, setting their resting activity to optimise hearing sensitivity at rest and in response to sound (**Hong et al., 2022**). During ageing or when the resting activity of adult IHC hair bundle is affected, cholinergic efferent axo-somatic synapses onto the IHCs reform for an unknown purpose, suggesting a reversion to the immature state **(Zachary and Fuchs, 2015; Lauer et al., 2012; Corns et al., 2018; Jeng et al., 2021**).

Recent data has suggested that these efferent fibres may arise from the iLOC system, though definitive confirmation is still required (**Corns et al., 2018; Jeng et al., 2021**). Importantly, if or how this cochlear rewiring influences the output of these neurons within the brainstem is not known. Understanding these shifts in biophysical properties is key to understanding how the hearing system might adapt to the ageing or dysfunctional cochlea.

In this study, we performed whole-cell recordings from iLOC neurons to investigate their biophysical characteristics throughout life. We found that iLOC neurons become more readily excitable with the onset of hearing, down-regulating their characteristic *I*_A_ current and up-regulating sodium channels and slow Kv2 channels in order to support bursting activity. In aged mice, iLOC neurons do not revert to an immature state and instead upregulate fast Kv3 subunits in order to support faster firing, resulting in an increased cochlear output. Utilising a mouse model of early IHC re-innervation via the post-natal hair cell specific deletion of *Myo7a* (**Corns et al., 2018**), we found that efferent re-innervation of the IHCs did not fully elicit premature ageing-like biophysics in iLOC neurons. Thus, iLOC neurons undergo distinct biophysical changes with age to underlie increased output to SGNs and therefore cochlear gain, and this appears to not be triggered solely by IHC dysfunction sufficient to elicit efferent re-innervation.

## Materials and Methods

### Sustainability statement

To promote environmental sustainability, the Hearing Group at the University of Sheffield utilizes resource-efficient laboratory workflows and holds a Silver LEAF accreditation. All research activities are conducted with a commitment to minimizing environmental impact. The laboratory adheres to sustainable practices, including strict “close the sash” protocols on fume hoods to optimise energy savings, the reduction of single-use plastics alongside the recycling of non-contaminated items, and the routine shutdown of equipment when not in use to reduce energy consumption. Material waste is minimized through the consolidation of reagent ordering to reduce shipping emissions, as well as the responsible segregation and recycling of non-hazardous lab plastics. Furthermore, computational analyses and imaging sessions are consolidated to minimize high-performance hardware runtime.

### Ethics statement

All the experiments were licenced under the Animals (Scientific Procedures) Act 1986 by the UK Home Office (PPL_ PP1481074). All animal procedures were performed in accordance with the University of Sheffield animal care committee’s regulations. Mice were kept in designated rooms with a 12-hour light/dark cycle, with Zeitgeber Time 0 (ZT0) corresponding to 06:00. Notably, all electrophysiology experiments were carried out with the mice being killed between ZT03 and ZT05. For ageing experiments mice were individually caged where necessary, usually following removal from a breeding pair at 6-9 months of age. For all brainstem experiments, animals were killed via an intraperitoneal overdose of pentobarbital (Dolethal; Vetoquinol, diluted to 100 mg/ml in saline). For cochlear immunolabelling, animals were killed via cervical dislocation followed by decapitation.

### Animals

All wild-type patch-clamp electrophysiology and immunolabelling experiments were performed on C57BL/6N mice bred in-house. This line has been utilised by other groups to investigate iLOC neurons (**Wu et al., 2020; Sitko et al., 2025**), and was selected for this due to their age-related hearing loss and IHC re-innervation evident at 12 months of age (**Jeng et al., 2021**).The *Myo7a^fl/fl^* animals were generated by the Mouse Genetics Programme at the Wellcome Trust Sanger Institute (Cambridge, UK). Exons 10 and 11 of Myo7a were flanked by loxP sites, and the *tm1c* allele was generated by crossing the *tm1a* line with FLPeR-expressing mice (Rosa26^Fki^). The experimental *tm1d* allele was obtained by crossing *tm1c* mice (*Myo7a^fl/fl^*) with Myo15-cre mice (kindly provided by Dr S. Safieddine; **Caberlotto et al., 2011**). For all experiments, *Myo7a^fl/fl^xMyo15-cre^+/-^*mice served as conditional knock-outs (KO) and *Myo7a^fl/fl^xMyo15-cre^+/+^* animals were used as littermate controls. Experiments were performed on both male and female animals without preference. For this work, pre-hearing iLOC neurons were those recorded from mice under post-natal day (P) 12 (<P12; P6-11), young adult was defined as 1-3 months old (1-3M; P24-81), and ageing animals were classed as any animal over a year old and subdivided into 12-14 months (12M; P366-412) or 18-20 months of age (18M; P557-596).

### Brainstem slicing

For electrophysiology experiments, immediately after decapitation the skin was pulled towards the nose such that the entire cranium caudal to the eyes was exposed. The skull was then placed into a beaker of pre-chilled and carbogen (95% O_2_, 5% CO_2_) equilibrated slicing solution, which itself had a layer of frozen slicing solution at the base to ensure rapid and continuous cooling. This solution varied depending on the age of the animal. Before post-natal day (P) 12 a classical sucrose base solution was utilised comprised of (in mM): 250 Sucrose, 2.5 KCl, 10 Glucose, 1.25 NaH_2_PO_4_, 26 NaHCO_3_, 0.5 L-ascorbic acid, 0.1 CaCl_2_, 4 MgCl_2_. Unless otherwise stated, for mice over P12 an N-Methyl-D-Glucamine (NMDG) based solution was utilised to ensure recordings could be carried out up to very old ages (**Ting et al., 2018**) and included (in mM): 93 NMDG, 2.5 KCl, 1.25 NaH_2_PO_4_, 30 NaHCO_3_, 20 HEPES, 25 Glucose, 5 L-ascorbic acid, 3 Myo-inositol, 3 Na-pyruvate, 10 MgCl_2_, 0.5 CaCl_2_, pH to 7.35 with HCl, osmolality ∼310 mOsm kg^-1^. After around 45 seconds of continuous mixing with the beaker sitting on ice, the skull was transferred to a petri dish containing a frozen base-layer of slicing solution with a small amount fresh and carbogen equilibrated solution on top to ensure effective cooling. In this solution, the brain was removed from the skull dorsally, ensuring continuous contact with the cooled solution wherever possible, and then place into 10 ml of fresh pre-chilled and carbogen equilibrated sucrose or NMDG slicing solution for transfer to the vibratome.

At the vibratome, the brain and slicing solution was poured onto a new petri dish also containing a base-layer of frozen slicing solution. The pia mater was then dissected away with fine forceps, and the brain was cut with a handheld vibratome blade at the upper part of the brainstem. The brainstem was then glued to the vibratome pedestal within the pre-cooled chamber surrounded by ice, and left to adhere for ∼45 seconds before filling with ice-cold slicing solution. With continuous carbogen bubbling, sections (100-150 µm) containing the lateral superior olive (LSO) were collected and temporarily kept within the sectioning chamber until sectioning was completed. Once all sections were collected, the subsequent recovery procedure varied depending on the slicing solution used. If a sucrose-based solution was utilised, samples were immediately transferred to a chamber containing aCSF (see below; Whole-cell electrophysiology) pre-warmed to 37°C and incubated in a water bath with continuous carbogen bubbling for 1 hour. Following this, the entire chamber was moved to room temperature and samples were left to cool and recover for at least one hour prior to the start of the experiment. For NMDG based solution, after slicing samples were immediately transferred into a chamber of NMDG slicing solution pre-warmed to 34°C and continuously bubbled with carbogen. As previously described (**Ting et al., 2018**), sodium was then gradually introduced to the slices in the 90ml chamber from a 2M stock of NaCl in the following manner: 0 min; 150 µl, 2 min: 150 µl, 4 min: 300 µl, 6 min: 600 µl, 8 min: 1200 µl, 10 min: transfer to carbogen equilibrated Hepes-holding solution, containing (in mM): 93 NaCl, 2.5 KCl, 1.25 NaH_2_PO_4_, 30 NaHCO_3_, 20 Hepes, 25 Glucose, 5 L-ascorbic acid, 3 Myo-inositol, 3 Na-Pyruvate, 2 MgCl_2_, 2 CaCl_2_, pH to 7.35 with NaOH, osmolality ∼310 mOsm Kg^-1^. Slices were left in Hepes-holding solution to recover at room temperature for at least 1 hour with continuous carbogen bubbling before experiments. Regardless of recovery procedure, slices were stored for a maximum time of 4 hours before being discarded.

### Whole-cell electrophysiology

Brainstem slices containing the LSO were transferred from the bubbled holding chamber into a microscope chamber containing carbogen equilibrated aCSF, comprised of (in mM): 125 NaCl, 2.5 KCl, 10 Glucose, 1.25 NaH_2_PO_4_, 26 NaHCO_3_, 2 Na-pyruvate, 3 Myo-inositol, 0.5 L-ascorbic acid, 1 MgCl_2_, 2 CaCl_2_, pH ∼7.4, osmolality ∼310 mOsm Kg^-1^. The slice was immobilised under a grid of dental floss attached to a stainless steel ring, and continually perfused via a peristaltic pump (Cole-Palmer, UK). The chamber was then placed into a custom rotating stage on an upright microscope (Olympus BX51), and the aCSF within the chamber was continually maintained at body temperature (35-37°C) with a combination of in-line solution heating and a thermometer controlled heated baseplate. Neurons were visualised via infrared illumination through Nomarski Differential Interference Contrast (DIC) optics through a 64x water immersion objective, captured through a monochrome camera (DMK41BU02.H; Imaging Source, Germany). Slices were maintained in this manner for a maximum of 1 hour, before being either processed for neuron tracing or discarded.

Recordings from neurons were carried out with soda glass capillaries pulled to a resistance of 3.3-3.6 MOhm (Narishige PP-830) with potassium gluconate intracellular solution and aCSF extracellular solution. Patch pipettes were manually coated with wax (Mr Zoggs SexWax, USA) to reduce pipette capacitance. For all whole-cell experiments, the intracellular solution was potassium gluconate based and contained (in mM): 97.5 K-gluconate, 32.5 KCl, 20 HEPES, 0.2 EGTA, 1 MgCl_2_, 2.5 Sodium phosphocreatine, 2 Na_2_ATP, 100 nM CaCl_2_, pH 7.2 with KOH, osmolality adjusted to ∼290 mOsm Kg^-1^. In experiments that required the tracing of recorded cells, biocytin (B4261, Merck, USA) at a concentration of 1 mg/ml was added to the intracellular solution on the day of use and sonicated at room temperature until fully dissolved. At the conclusion of recording a neuron with biocytin intracellular solution, the patch pipette was slowly retracted such that a giga-seal was reformed to prevent excessive cell damage and loss of intracellular biocytin. Following this, the slice was removed from the recording chamber and immediately placed directly into 1 ml of 4% paraformaldehyde (PFA) in phosphate buffered saline (PBS) for overnight fixation. For cell attached recordings, the intracellular solution mimicked the aCSF as close as possible and contained (in mM): 125 NaCl, 2.5 KCl, 10 Glucose, 10 HEPES, 1 MgCl_2_, 2 CaCl_2_, pH to 7.4 with NaOH and osmolality adjusted to ∼290 mOsm Kg^-1^ with sucrose.

For experiments which included pharmacologically active compounds, a perfusion system was utilised to apply drug directly onto the patched cell. TEA-chloride (86615, Sigma-Aldrich, USA) was readily soluble, and dissolved directly into aCSF. RY785 was dissolved in DMSO to create a 5 mM stock solution, which was diluted 5000x in aCSF to create a working solution of 2.5 µM. As such, for RY785, the control aCSF was supplemented with DMSO to ensure concentrations of the organic solvent matched. Recordings were taken at least 2 minutes after the onset of drug perfusion, or when the effect had reached an evident steady state.

Data acquisition was carried out with pClamp software utilising a Digidata 1440A digitiser (Molecular Devices, USA). Recordings were sampled at 5 kHz and low pass filtered (8-pole Bessel) at 2.5 kHz for potassium current protocols or sampled at 20 kHz and low pass filtered at 5 kHz for sodium current protocols. Recordings were analysed via Clampfit (Molecular Devices, USA) and Origin 2021 (OriginLab, USA). In voltage clamp protocols, the membrane potentials were corrected for the residual series resistance *R_s_*, which was actively compensated at 60% for all recordings, and a measured liquid junctional potential (LJP) of +8 mV between the aCSF and potassium gluconate intracellular solution. All voltage clamp protocols were carried out from a holding potential of -70 mV, and therefore -62 mV following LJP correction.

### Brainstem immunolabelling and biocytin tracing

In order to collect samples for immunolabelling experiments, animals were culled as outlined above, and each brain was dissected out at room temperature and place immediately into 2.5 ml of 4% PFA in PBS. Samples were left to fix overnight at 4°C, and transferred to PBS the following day if being used immediately or into PBS supplemented with 5mM sodium azide for long term storage. Sectioning on the vibratome was carried out at room temperature in a chamber filled with PBS.

PFA fixed brainstem slices containing the LSO, were first washed in PBS three times to remove residual PFS. Slices were then placed into a blocking/permeabilization solution of 1% Triton X-100 with 5% normal horse serum (NHS) at room temperature for two hours. Brainstem slices were then immunolabelled overnight at 37°C with goat-IgG anti-ChAT (Dilution 1:500, AB144P, Merck Millipore, USA) diluted in 1% Triton X-100 with 1% NHS. Slices were washed with PBS three times for 15 minutes each, and then placed into a secondary antibody solution of donkey anti-goat 557 (Dilution 1:1000, NL001, Bio-Techne, USA) and streptavidin 647 (Diluted 1:1000, 521374, Invitrogen, USA) diluted in 1% Triton X-100 with 1% NHS and incubated at 37°C for 2 hours. Slices were then washed three times in PBS as before and immediately mounted onto cavity microscope slides (N/A114, ACADEMY, UK) in VECTASHIELD anti-fade mounting medium (H-1000, Vector Laboratories, USA) and sealed with nail varnish. Image stacks of the entire LSO were acquired on a Nikon W1 Spinning Disc Microscope via a 20x Plan Apochromat Lambda D objective (Wolfson Microscopy Facility at the University of Sheffield). Resulting image stacks were analysed and processed with Fiji ImageJ software. For biocytin facilitated measurement of the tonotopic position the tonotopic axis was drawn manually by an investigator blinded to age, as immunolabelled samples were numbered sequentially without reference to age of genotype, and from this the tonotopic position of the traced cell as a percentage (where 0% is the lowest frequency and 100% is the highest) was estimated.

### Cochlea immunolabelling

Inner ears were dissected out of the skull and a small incision was made into the apical cap of the cochlea. Inner ears were then perfused with a fixation solution of 4% PFA in PBS through the round and oval windows using a 100 µl pipette tip. After around 30 seconds of perfusion, inner ears were then placed into a 1.5 ml Eppendorf tube containing PFA solution at room temperature for a further 20 minutes. Inner ears were then washed three times for 5 minutes each in PBS, and if to be kept for long term storage were transferred to a solution of PBS with 5 mM sodium azide. The apical turns of the cochleae were then dissected out using forceps, washed in PBS, and incubated with a blocking solution of 5% NHS and 0.5% Triton X-11 in PBS at room temperature for one hour. The samples were then immunolabelled with primary antibodies diluted in 1% NHS and 0.5% Triton X-100 in PBS overnight at 37°C. Primary antibodies used were mouse-IgG1 anti-Myo7a (Diluted 1:100, 138-1-c, Proteus Biosciences, USA), mouse IgG2a anti-class III beta-tubulin (Diluted 1:1000, 801201, Biolegend, USA), rabbit IgG anti-SK2 (Diluted 1:100, P0483, Sigma Aldrich, USA). Following incubation with primary antibodies, samples were washed on a shaker three times in PBS for 5 minutes per wash. The cochlear turns were then incubated with species appropriate Alexa Fluor or Northern Lights fluorescently conjugated secondary antibodies diluted in 1% NHS and 0.5% Triton X-100 in PBS for 1 hour at 37°C. Following another round of three 5 minute PBS washes, samples were then mounted under glass coverslips using VECTASHIELD anti-fade mounting medium (H-1000, Vector Laboratories, USA) and sealed with nail varnish. Images from the same approximate region of the cochlear spiral were acquired using a Zeiss LSM 880 Airyscan microscope utilising a Plan-Apochromat 63x Oil DIC M27 objective at the Wolfson Microscopy Facility (University of Sheffield). Image stacks were processed with Fiji ImageJ software.

### Statistical analyses and experimental design

The experimental unit for this work was defined as a single cell, and a maximum of 4 cells were collected from each mouse to avoid pseudoreplication. Experimental units vary between experiments, and as such both animal and cell n-numbers are reported within the figures and their associated figure legends. All recordings were made within the visible boundary of the lateral superior olive (LSO), and any cell with a series resistance higher than 11 MOhm was discarded. All datasets were tested for normality with a Shapiro-Wilk test. If data was normally distributed in all groups, statistical comparisons were made with a Student’s t-test for individual comparisons or by analysis of variance (one- or two-way ANOVA followed by Tukey’s post-test) for multiple comparisons. If any groups were not normally distributed, statistical comparisons were instead carried out with a Mann-Whitney U-test or Kruskal-Wallis ANOVA with Dunn’s post-test. For tonotopy analysis, linear regression analysis was carried out with Bonferronni’s correction for multiple comparisons. All statistical analyses were carried out with OriginPro (OriginLabs, USA). *P* < 0.05 was defined as the threshold for statistical significance, and the exact *P* value for all statistically significant results are shown with the data. All normally distributed graphical data shown is plotted as the mean ± standard deviation, and all non-normally distributed data is plotted as median ± 10/90^th^ percentile.

## Results

### Identification of iLOC neurons within the LSO

To investigate the biophysical characteristics of iLOC neurons throughout life whole-cell patch-clamp electrophysiology was utilised, with the vast majority of patching experiments carried out with a biocytin containing intracellular solution to allow patched cells to be traced (**Figure 1A-D**). The identity of iLOC neurons was confirmed if they were constrained within the boundaries of the LSO (**Figure 1A)** and expressed choline acetyl-transferase (ChAT; **Figure 1B-D)**. Biophysically, iLOC neurons could be reliably differentiated from other neurons within the LSO by a pronounced delay to the first spike following small current injections (**Figure 1E**), as previously described (**Sterenborg et al., 2010**). In voltage-clamp configuration, iLOC neurons also lacked the inward H-type current present during hyperpolarising voltage steps (*I*_H_) showing only a leak current (**Figure 1F**), and exhibited a prominent inactivating outward A-type potassium current (*I*_A_) (**Figure 1G) (Fujino et al., 1997; Sterenborg et al., 2010**) which is carried by Kv4 potassium channel subunits (**Birnbaum et al., 2004; Leijon and Magnusson, 2014**). By stepping the pre-conditioning pulse from -82 to +22 mV the *I*_A_ could be progressively inactivated (**Figure 1H**), and the size of this inactivating current could be plotted and fitted with a Boltzmann equation (**Figure 1I**) (**Bekkers, 2000**). This approach permitted accurate measurement of current size and half-inactivation voltage without conflation with other cellular potassium currents.

**Figure 1.**
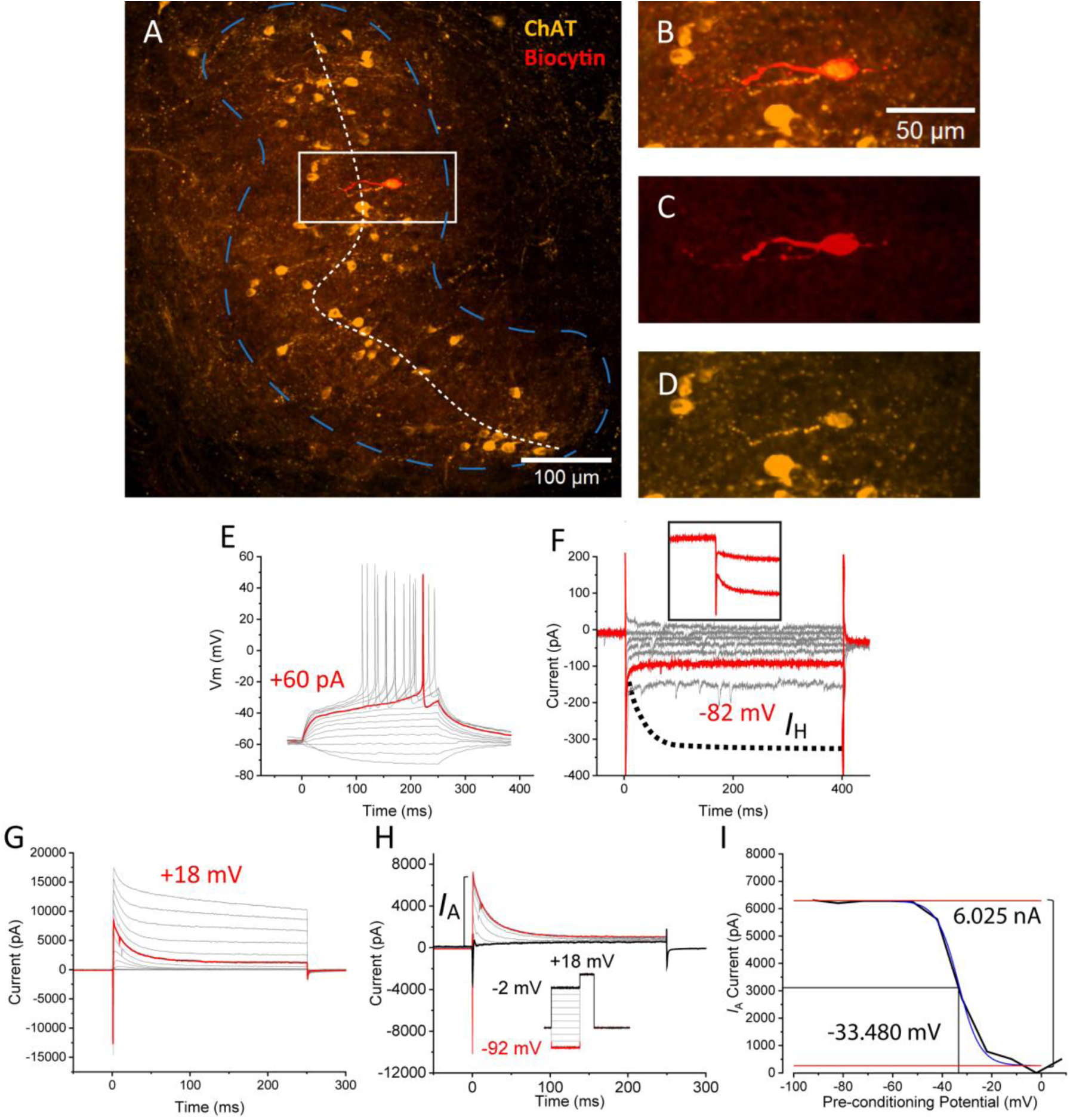
Identification of intrinsic lateral olivocochlear neurons. *A*, Example coronal section of the P6 C57BL6/N brainstem containing the lateral superior olive (LSO; bordered in blue), with the ChAT positive (orange) intrinsic lateral olivocochlear (iLOC) neurons evident running throughout, projecting dendrites perpendicular to the tonotopic axis (dashed white line). The white square highlights a biocytin filled iLOC neuron fluorescently labelled with fluorophore conjugated streptavidin (Red). ***B****-**D***, Magnified image of the biocytin labelled iLOC neurons in ***A***, possessing both biocytin (***C***) and expression of ChAT (***D***). ***E,*** Verified iLOC neurons possessed the characteristic delay to first spike following near threshold current injections and no visible *I*_H_ sag in response to hyperpolarising current steps from -20 pA to +100 pA. ***F*,** iLOC neurons also do not show the inward *I*_H_ in response to hyperpolarising voltage steps in 10 mV intervals from -32 to -92 mV from a conditioning potential of -42 mV. For comparison, the inset in ***F*** shows the inward *I*_H_ current recorded from a principle neuron of the LSO, and the dotted line on the main panel shows how the inward current may look is *I*_H_ was present. ***G***, iLOCs also possessed a prevalent rapid inactivating A-type current (*I*_A_) in response to positive voltage steps in intervals of 10 mV from a pre-conditioning step of -82 mV. ***H***, *I*_A_ can be inactivated by applying pre-conditioning voltage steps between -92 (Red) to +38 mV (-2 mV example trace in black, with only -92 to -2 steps shown in the panel for clarity) followed by a test potential of +18 mV, which allows for an accurate measurement of current size. ***I***, By plotting the pre-conditioning potential against the size of the inactivating current, measured by subtracting the steady state value from the peak, the size of *I*_A_ as well as the half-inactivation value (6.025 nA and -33.480 mV respectively in this example) could be calculated.

### Cell size and the biophysical characteristics of potassium currents in iLOC neurons change with age

To characterise how iLOC biophysics adapt during development and then determine how this compares with ageing, whole-cell patch clamp experiments were conducted on the iLOC neurons of wild-type C57BL/6N (6N) mice. From these experiments, the inactivating *I*_A_ was particularly evident in pre-hearing animals in response to voltage steps from a -62 mV to +68 mV in 10 mV increments from a -82 mV holding voltage (**Figure 2A**) compared to older ages (**Figure 2B-D**). Cellular capacitance, which provides a measure of cell surface area, decreased after the onset of hearing and then remained stable throughout the following life-course (**Figure 2E**)^a^.

**Figure 2.**
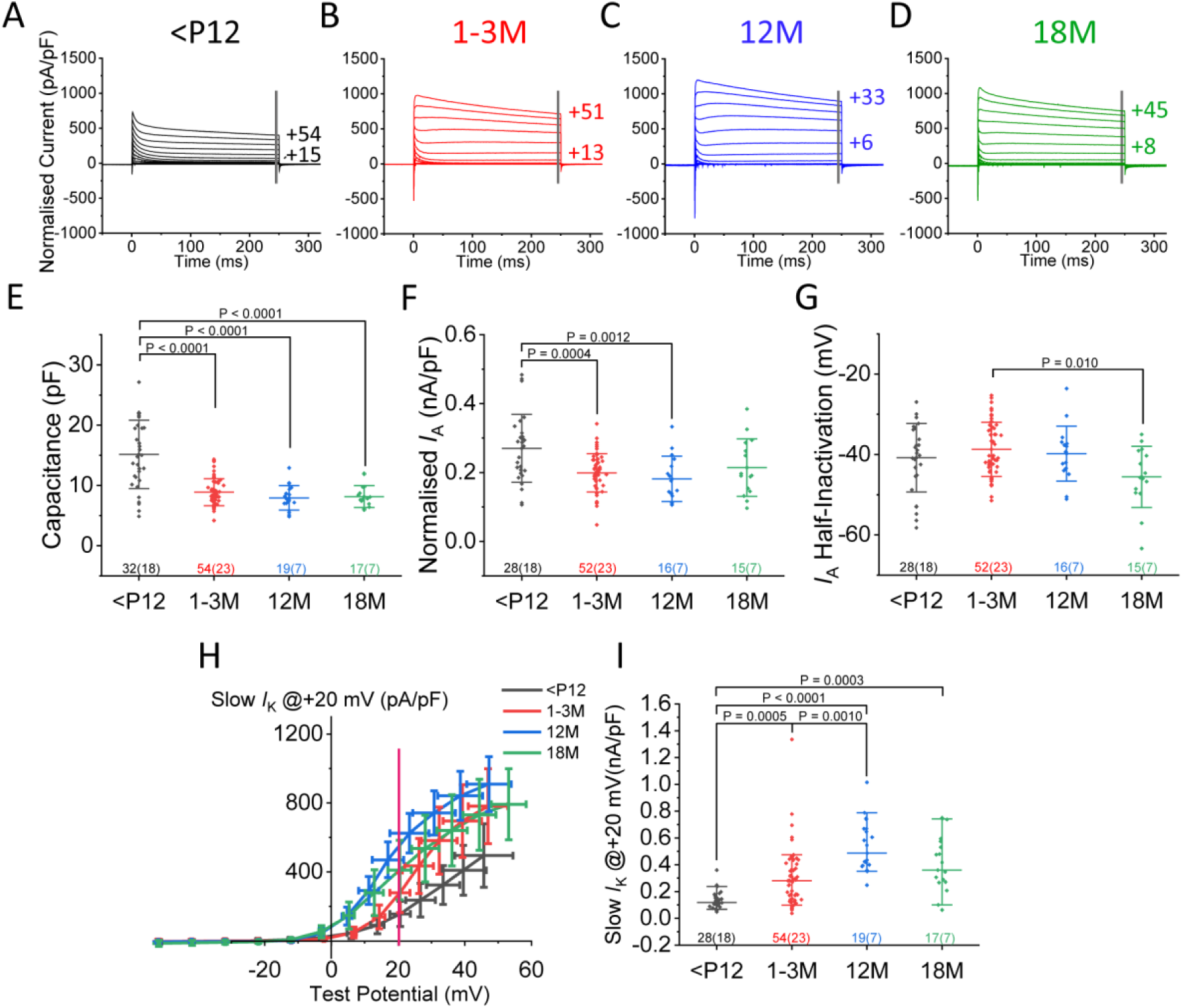
Changes in cell surface area and membrane currents in lateral olivocochlear neurons at different ages. ***A****-**D***, Average total current recordings from iLOC neurons of pre-hearing (***A*** *-* <P12; P6-P11, n=32), young adult (***B*** – 1-3M; P24-P89, n=54), 12 months (***C*** – 12M; P366-P416, n=19), and 18 months old (***D*** – 18M; P557-P564, n=16) mice. Current responses were obtained by stepping the voltage from -62 mV to +68 mV in 10 mV increments from a holding potential of -82 mV. ***E***, Membrane capacitance of iLOC neurons as a function of age. ***F***, Peak amplitude of the isolated A-type outward potassium current (*I*_A_) measured via stepwise pre-conditioning inactivation from -92 mV to +38 mV and a test pulse of +18 mV. ***G***, Half-inactivation of *I*_A_ obtained via the methodology outlined in Figure 1I. ***H***, Size of the non-inactivating outward current, measured from the end of the traces as shown in ***A-D*** (gray line) plotted against the test potential. ***I,*** The steady-state outward potassium current measured at +20 mV as shown by the magenta line in ***H***. Statistics were performed with a one-way ANOVA followed by Tuckey’s post-test or a Kruskal-Wallis ANOVA followed by Dunn’s post-test, with exact *P* values indicated over each statistically significant comparison. The number of iLOC neurons and mice are shown beneath the data in the format; iLOC neurons (mice).

The size of *I*_A_, calculated from the current size normalised to cell capacitance, also showed a reduction after the onset of hearing, and did not significantly reduce further in ageing mice of either group relative to young-adults (**Figure 2F**)^b^. Whilst the size of *I*_A_ decreased in young adult iLOC neurons relative to pre-hearing neurons, its half-inactivation characteristics remained stable up until one year of age, becoming significantly decreased only at the oldest age tested of 18 months relative to young-adult animals (**Figure 2G**)^c^. The total steady-state potassium current in iLOC neurons (slow *I*_K_), which was measured at the end of the traces shown in **Figure 2A-D**, increased with age (**Figure 2H**). Measuring this current at +20 mV indicated a significant increase with the onset of hearing and ageing that peaked at around 12 months (**Figure 2I**)^d^. 20 mV was selected as the voltage for comparison as it captures the opening of higher voltage potassium channels (**Figure 2H**), but the effect of residual series resistance is minimal, so values are not lost due to failure to reach the true voltage. Notably, further studies will utilise higher voltage steps, as this would permit the fitting of a Boltzmann to the traces shown in **Figure 2H** and permit analysis of any shape differences. Taken together, these results suggest that both the inactivating *I*_A_ and the total steady-state outward potassium current in iLOC neurons change before and after hearing onset, and the steady-state outward current grows further as iLOC neurons age up until one year, rather than reverting to more immature values as animals age beyond young-adulthood. As such, this data does not align with the original hypothesis that ageing iLOC neurons would revert to an immature biophysical profile.

### The rapid component of the non-A type outward current and the sodium current in iLOC neurons are differentially regulated with age

In order to investigate iLOC membrane currents throughout life without interference from *I*_A_, a holding step of -32 mV was utilised to voltage-inactivate *I*_A_, removing the distinctive “saw-tooth” component from the total outward potassium current (**Figure 3A-D**). Using these recordings, the rapid component of the outward current was measured at 2 ms after the onset of the voltage step (**Figure 3E**). In contrast to the steady-state outward current, when measured at +20 mV the fast non-*I*_A_ outward potassium current remained stable before and after the onset of hearing, and rose as the animals aged between 1-3 and 12 months, remaining significantly elevated in 18 month old aged mice relative to pre-hearing and young adult animals (**Figure 3F**)^e^. The sodium current in iLOC was investigated using this same -32 mV holding protocol, as the inactivation of *I*_A_ made it possible to record the inward sodium transient with minimal contamination from this fast-activating outward current (**Figure 3G**) (**Milescu et al., 2010**). Using this methodology, the size of *I*_Na_ increased during the onset of hearing, remaining stable between young adult and 12 month aged animals, and then decreased between 12 and 18 months of age (**Figure 3H**)^f^. The potential at which the sodium current peaked also shifted to be more negative as iLOC neurons aged at 12 and 18 months relative to both pre-hearing and young-adult animals, but was unaffected before and after the onset of hearing (**Figure 3I**)^g^.

**Figure 3.**
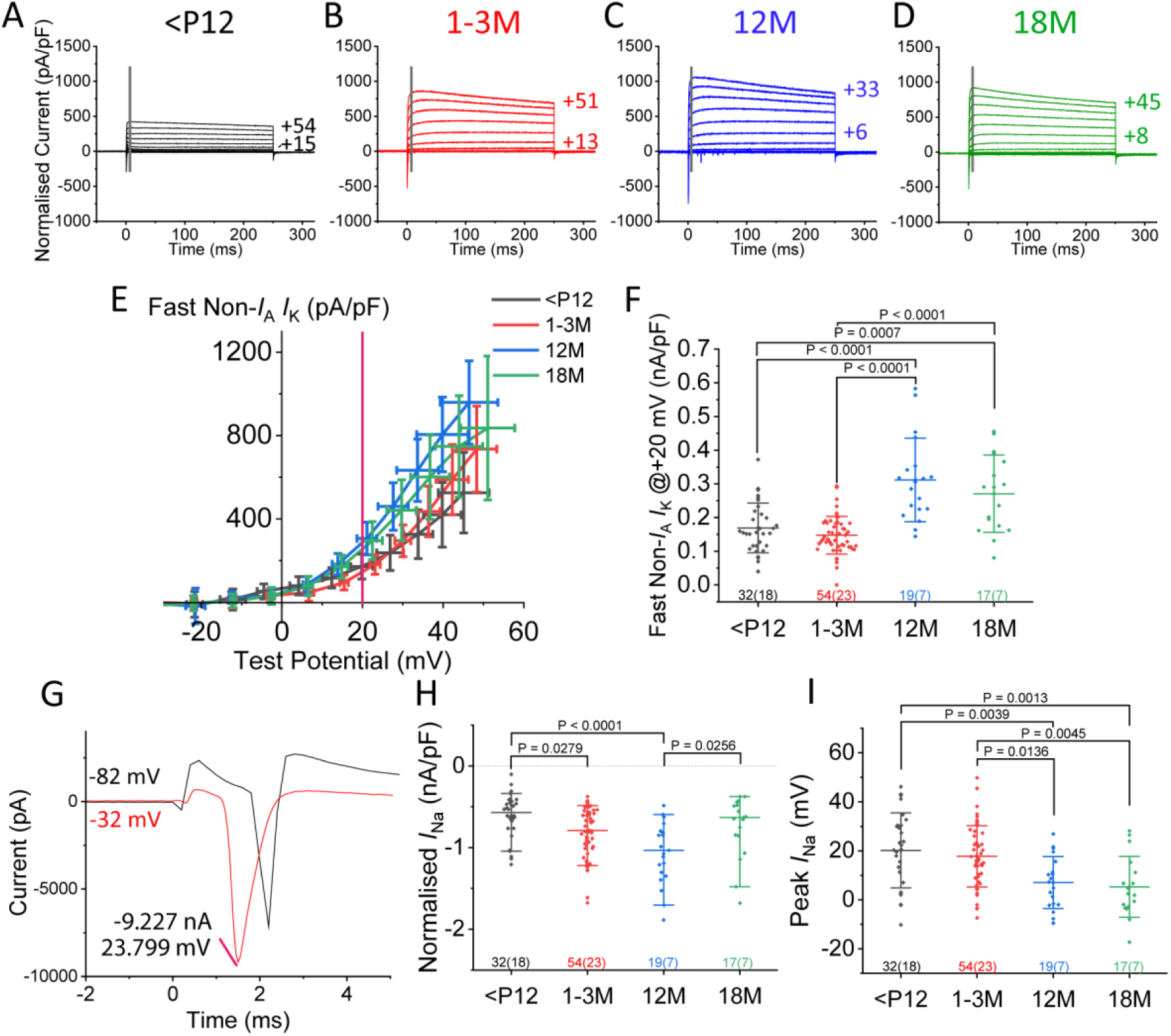
The non-*I*_A_ fast potassium current and sodium current shift in size and kinetics with age. ***A-D***, Average total current recordings from iLOC neurons of pre-hearing (***A***. <P12; P6-P11, n=32 cells from 18 animals), young adult (***B***, 1-3M; P24-P89, n=54 cells from 23 animals), 12 months (***C***, 12M; P366-P416, n=19 cells from 7 animals), and 18 months old (***D***, 18M; P557-P564, n=13 cells from 6 animals) mice. Current responses were obtained by stepping the voltage from -62 mV to +68 mV in 10 mV increments from a holding potential of -32 mV to inactivate *I*_A_. ***E***, Size of the fast non-*I*_A_ outward potassium currents measured at 2 ms after the onset of the voltage step as shown in ***A-D*** (gray line) plotted against the test potential. ***F,*** The density of the fast outward potassium current measured at +20 mV (magenta line in ***E***). ***G,*** Example inactivation of *I*_A_ utilising a -32 mV holding potential (Red) to permit sodium current measurements with less interference from the fast *I*_A_ than the standard -82 mV protocol (Black). In this example, generated from 10 mV voltage steps and states actual voltage after correcting for residual series resistance, the sodium peak was 9.227 nA at 23.799 mV from a voltage step of -12 mV. ***H***, Peak of the sodium current (*I*_Na_) in iLOC neurons obtained by stepping the traces outlined in ***A-D***. Current size was normalised to capacitance. ***I,*** Voltage at which *I*_Na_ peaked, measured as shown in ***G***. Statistics were performed with a one-way ANOVA followed by Tuckey’s post-test or a Kruskal-Wallis ANOVA followed by Dunn’s post-test, with exact *P* values indicated over each statistically significant comparison. The number of iLOC neurons and mice are shown beneath the data in the format; iLOC neurons (mice).

**Figure 3.1.**
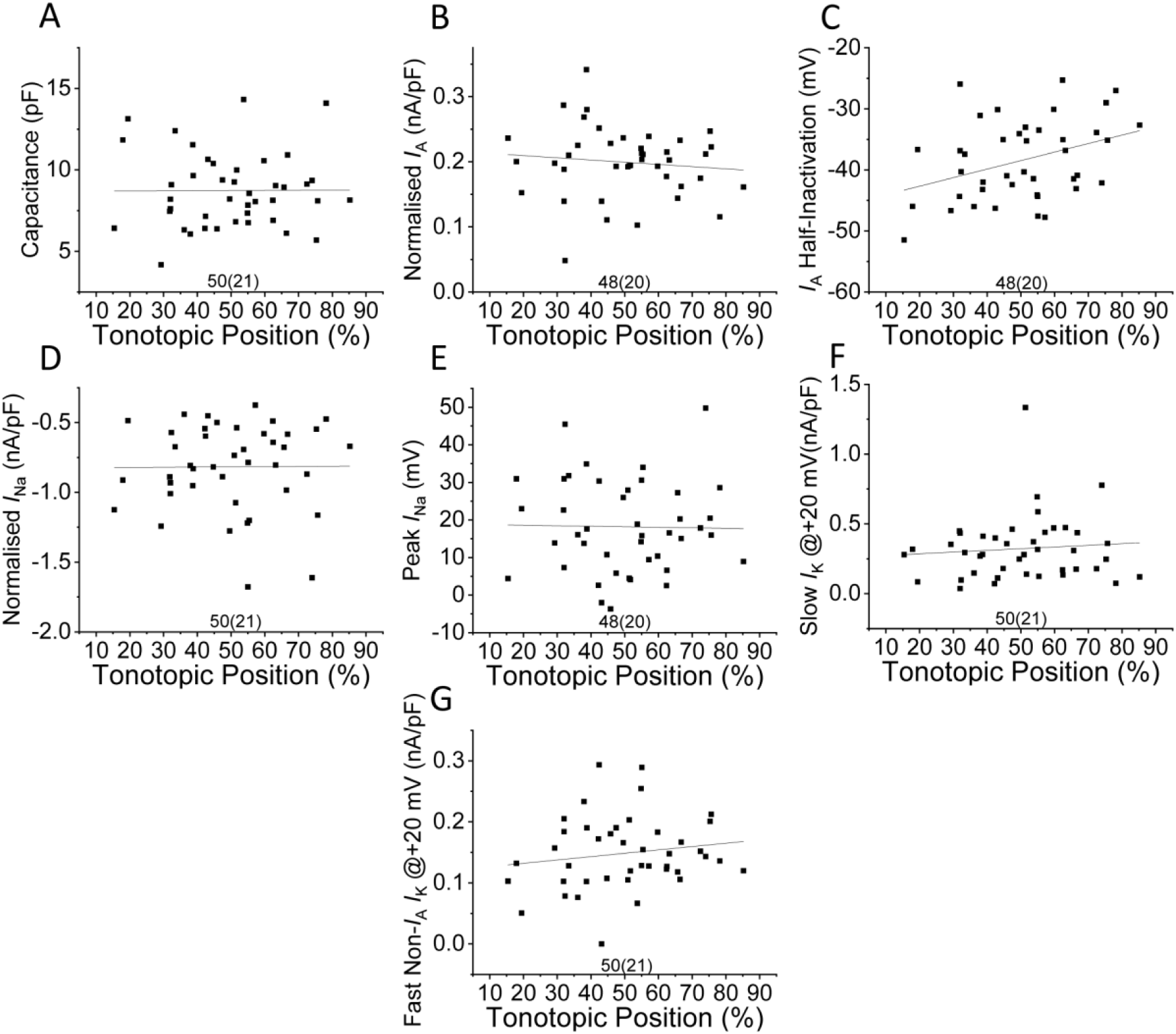
Relationship between voltage clamp metrics and tonotopy. Extended data for Figure 3. ***A-G***, iLOC neuron metrics from mature animals relative to tonotopic position of the recorded cell, including capacitance (***A***), *I*_A_ density (***B***), *I*_A_ half-inactivation voltage (***C***), *I*_Na_ density (***D***), voltage of peak *I*_Na_ (***E***), size of the slow (***F***) and fast (***G***) non-*I*_A_ outward potassium channel. 0% indicates the most lateral and low frequency part of the LSO and 100% indicates the most medial and highest frequency. The number of iLOC neurons and mice are shown beneath the data in the format; iLOC neurons (mice). Statistical analysis was conducted with a linear regression analysis in OriginPro. As none of the findings are statistically significant following Bonferronni’s correction for multiple comparisons (Alpha = 0.0071), exact *P* values are not shown.

**Figure 3.2.**
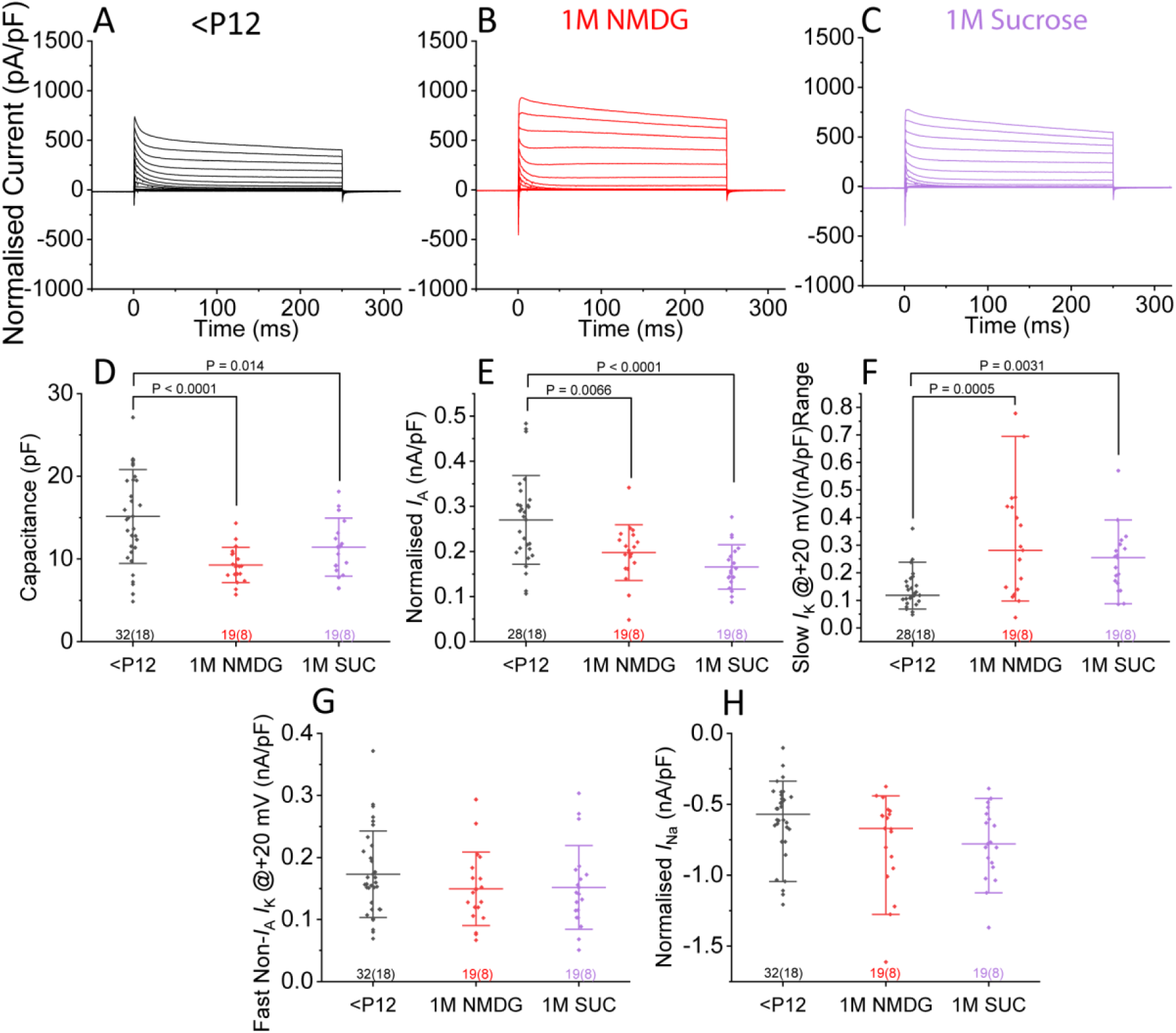
The NMDG slicing solution does not induce the biophysical changes seen between pre- and post-hearing iLOC neurons. Extended data for Figure 3. ***A-C***, Average total current recordings of pre-hearing (***A*** – <P12; P6-P11, n=32) iLOC neurons prepared with sucrose solution, alongside young adult iLOC neurons prepared with an NMDG based slicing solution (***B*** – 1M NMDG; P25-34, n=19) or a sucrose based slicing solution (***C*** – 1M Sucrose; P24-35, n=19). Current responses were obtained by stepping the voltage from -62 to +68 mV in 10 mV increments. ***E-H,*** As described in Figure 2 and **3**, values were obtained for membrane capacitance (***D***), *I*_A_ density (***E***), and the steady-state potassium current (***F***) measured from the end of the traces shown in ***A-C***. ***G***, Density of the fast non-inactivating outward current measured from 2 ms after the onset of the voltage step after a -32 mV holding step and compared at +20 mV. ***H,*** Size of *I*_Na_ when normalised to membrane capacitance. Statistics were performed with a one-way ANOVA followed by Tuckey’s post-test or a Kruskal-Wallis ANOVA followed by Dunn’s post-test, with exact *P* values indicated over each comparison. The number of iLOC neurons and mice are shown beneath the data in the format; iLOC neurons (mice).

Utilising a similar approach as above, the inward leak current of iLOC neurons were also investigated with hyperpolarising voltage steps from -32 to -92 mV from a holding of -42 mV. Using this protocol, steady-state inward currents in iLOC neuron responses were small, and when normalized to cell capacitance averaged ∼10 pA/pF at -92 mV at the end of the voltage step, a value that remained statistically unchanged at every age assessed (*P* = 0.0977. <P12; n = 18, median = -8.135 pA/pF IQR: [-11.572, -4.849 pA/pF]. 1-3M; n = 41, median = -7.144 pA/pF IQR: [-16.299, -3.942 pA/pF]. 12M; n = 9, median = -10.488 pA/pF IQR: [-11.506, - 5.956 pA/pF]. 18M; n = 5, median = -8.990 pA/pF IQR: [-12.190, -5.688 pA/pF]. Statistics were performed with a Kruskal-Wallis ANOVA followed by Dunn’s post-test.)^h^.

In order to determine whether any tonotopicity existed in the biophysical characteristics investigated, values were plotted against tonotopic position for young-adult iLOC neurons (**Figure 3.1**)^i^. Utilising linear regression analysis, it was revealed that none of the biophysical characteristics showed any correlation with tonotopic position following correction for multiple comparisons. Of note, capacitance, input resistance, and resting membrane potential have been reported by other groups to have no tonotopic gradient, which aligns with these findings (**Sterenborg et al., 2010**).

To carry out high-quality whole-cell recordings from pre-hearing up to such advanced ages, it was necessary to employ an NMDG based slicing solution for neurons over the age of P12 (**Ting et al., 2018**), whereas a more simple sucrose solution was employed for younger animals as previously done (**Richardson et al., 2022**). However, utilising NMDG slicing solutions for young pre-hearing neurons, as reported by other groups, prohibited high quality seal formation and thus usable current clamp recordings (**Ting et al., 2014**). To ensure the different solution were not inducing changes in iLOC biophysics, some of the P24-32 iLOC neurons were patched with the sucrose slicing solution and compared with age-matched neurons patched with the NMDG slicing solution used for the adult and ageing experiments (**Figure 3.2**). Notably, iLOC neurons patched with NMDG-based slicing solution retained their biophysics, including capacitance, *I*_A_ and *I*_Na_ current densities, and the slow and fast potassium current densities (**Figure 3.2D-H**). Indeed, despite the smaller dataset, the decrease in cell capacitance and *I*_A_ current size, as well as the increase in the slow potassium current, were still significantly shifted between pre-hearing and mature iLOC neurons utilising the sucrose-based slicing solution (**Figure 3.2D-F**)^j-l^. No statistically significant difference was found in the size of the fast non-*I*_A_ potassium current, in line with the preceding data (**Figure 3.2G)**^m^. The absence of a significant upregulation of the sodium current in this comparison is likely due to the more restrictive age-grouping to ensure sucrose and NMDG groups are closely matched, resulting in fewer included iLOC neurons in the NMDG group and a reduced statistical power **Figure 3.2H**)^n^.

### The non-A type components of mature outward potassium currents in iLOC neurons are carried by Kv3 and Kv2 channels

To identify the ion channels underpinning the non-inactivating outward potassium current, to better understand the changes occurring with the onset of hearing and ageing, we followed a previously described pharmacological approach outlined by **Johnston et al., (2010)**. Given the high voltage threshold of these non-inactivating currents, and the speed at which the rapid component activates which is measured at 2 ms, it was hypothesized that Kv3 comprised at least a proportion of this outward current. To test this, we utilised 1 mM tetraethylammonium chloride (TEA) which, despite being a non-selective potassium channel blocker, acts as a relatively selective blocker of Kv3 channels at these low concentrations (**Johnston et al., 2010**). Notably, this concentration is also sufficient to block BK and Kv7 potassium channels (**Johnston et al., 2010**), however, the high threshold (∼10 mV; **Figure 3E**) and fast activation of the investigated current are more indicative of Kv3 channels, and more detailed follow-up investigations are required to definitively rule out a role any role of BK or Kv7 in iLOC neurons. At 1 mM, TEA could be seen to have a clear effect on the net outward current as shown by the example traces in **Figure 4A-B**, and in line with the literature (**Leijon and Magnusson, 2014**). For the rapidly activating outward component measured at 2 ms after the onset of stimulus, the blockade of Kv3 more greatly reduced the early response to the voltage step relative to the steady-state component (**Figure 4C**). Due to the time taken for drugs to equilibrate in the bath, which was typically over the course of minutes, raw measurement of the fast and slow current densities proved unreliable due to progressive current run-down (**Numann et al., 1987**). However, the effect of 1 mM TEA on the fast component and recovery after the wash could be visualised by quantifying the ratio between the fast and steady-state current sizes. Using this method, 1 mM TEA clearly reduced the size of the fast potassium current within iLOC neurons relative to the lesser affected slow component, and this effect could be reversed following washing with aCSF (**Figure 4D**)°.

**Figure 4.**
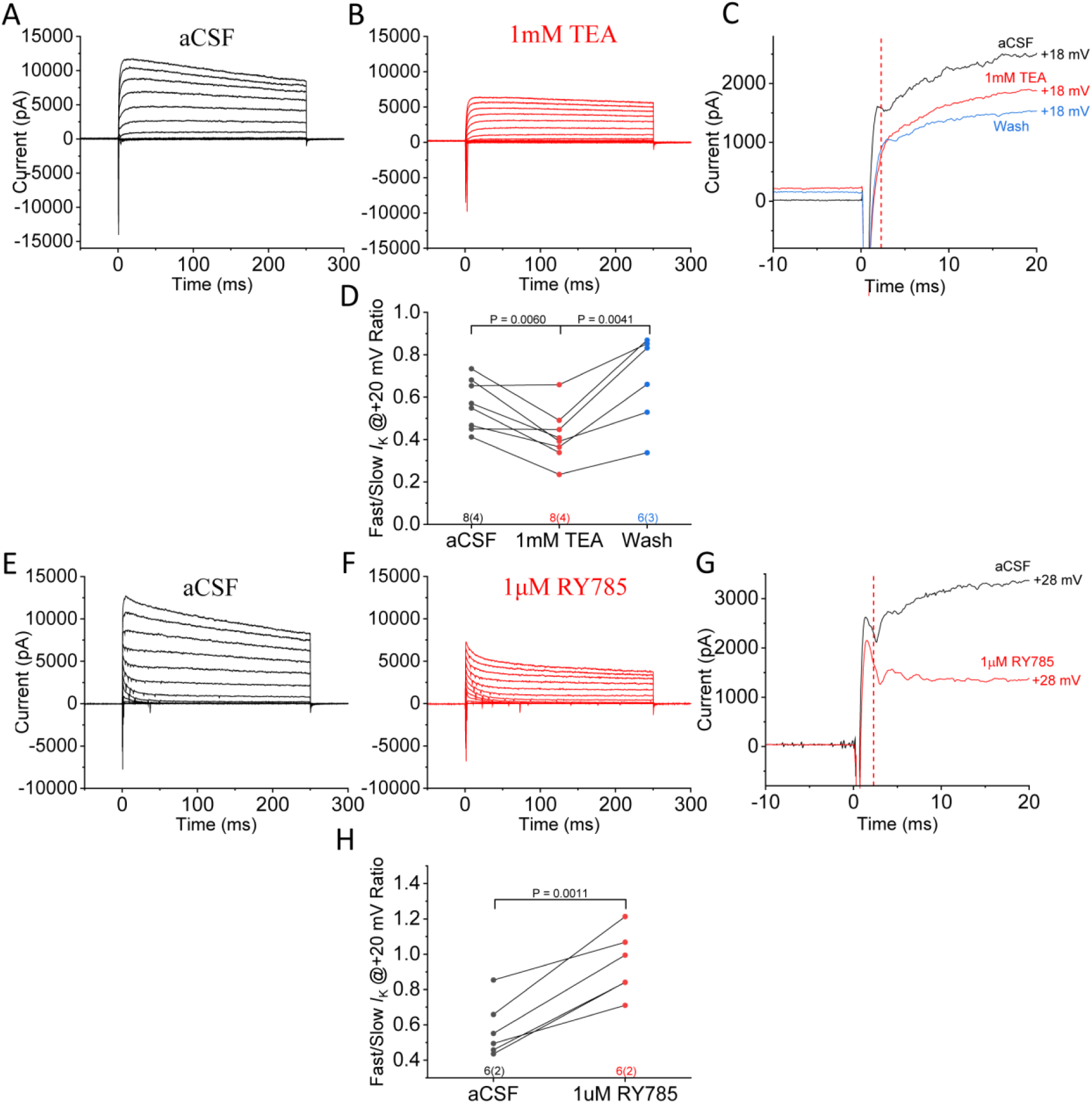
Mature iLOC neurons possess Kv3 and Kv2 potassium currents. ***A***-***B***, Example current recordings of a P26-P35 iLOC neuron before (***A***) and after (***B***) the application of 1 mM TEA, in response to 10 mV voltage steps between -62 mV and +68 mV after *I*_A_ is conditionally deactivated with a -32 mV pre-conditioning pulse. ***C***, Comparison of traces ***A*** and ***B*** zoomed into the first 20 ms of the +18 mV test pulse, with the red dotted line indicating 2 ms after the onset of the test pulse. ***D***, Ratio calculated as the fast potassium current (measured at 2 ms) divided by the steady state at +20 mV during the stages of drug application. ***E-F***, Example recordings from an iLOC neuron before (***E***) and after (***F***) application of 1 µM RY785 in response to 10 mV voltage steps between -62 mV and +68 mV, with *I*_A_ present due to a pre-conditioning pulse of -82 mV to highlight shape changes. ***G***, Comparison of before and after RY785 drug application at a test pulse of +28 mV following a -32 mV pre-conditioning step to inactivate *I*_A,_ with the red dotted line representing 2 ms after the onset of the test pulse. ***H***, Ratio between the fast (2 ms) and steady state potassium currents before and after drug application measured at +20 mV. Due to missing values, statistics were performed with paired-sample t-tests followed by suitable Bonferroni correction for multiple comparisons (***D***) or a two-sample t-test (***H***) with exact corrected *P* values indicated over each comparison. The number of iLOC neurons and mice are shown beneath the data in the format; iLOC neurons (mice).

With Kv3 identified as being a significant carrier of the fast component, we next investigated the slower outward potassium current of iLOC neurons. As this steady-state current also opened at a high threshold, and appears to have slower activation kinetics than Kv3 (**Figure 4C**), we hypothesized that Kv2 subunits may be present in iLOC neurons (**Johnston et al., 2010**). To test this hypothesis, we utilised the specific Kv2 family blocker RY785 at a concentration of 1 µM. Strikingly, when applying test pulses from a holding potential of -82 mV, application of RY785 caused iLOC neurons to adopt an outward current profile (**Figure 4E-F**) reminiscent of that in pre-hearing neurons (**Figure 2A**). Utilising an *I*_A_ inactivating protocol with a -32 mV pre-conditioning step, the application of RY785 had little effect on the rapid component at 2 ms, but drastically reduced the size of the steady-state potassium current (**Figure 4G-H**)^p^. Notably, further pharmacological experiments are required to determine the composition of these channels, whether they are Kv2 homomers or heteromers containing other subunits. Taken together, this data indicates that alongside Kv4 subunits (**Leijon and Magnusson, 2014**), both Kv2 and Kv3 are major contributors to the outward current seen in post-hearing iLOC neurons.

### The onset of hearing and ageing have distinct effects on the spiking characteristics of iLOC neurons

To determine the functional consequences elicited by the shifts in iLOC potassium biophysics with age, spiking activity was investigated in response to step-wise 10 pA depolarising current injections (**Figure 5A-H**). From these recordings, values were obtained for the iLOC resting membrane potential measured as the average value from traces during the first 20 ms of the recording prior to stimulus onset, which only significantly decreased in 1-3 month old animals relative to pre-hearing neurons, with no other statistically significant differences between the age groups (**Figure 5I**)^q^. Further experimentation with a greater number of cells in the ageing groups is required to determine if this is an artefact of statistical power or whether resting membrane potential trends towards more depolarised values with ageing. When normalised to the current threshold, the most striking change in action potential metrics with age was the progressive loss of the characteristic delay to the first spike in response to current injection (**Figure 5J**). Due to missing values and the non-parametric nature of the data, global analysis of the data shown in **Figure 5J** could not be carried out, however the delay to the first spike at +50 pA was significantly decreased in ageing animals of both groups relative to pre-hearing neurons (*P* = 0.0023. <P12; n = 24, median = 48.00 ms IQR: [13.25, 68.28 ms]. 1-3M; n = 62, median = 18.90 ms IQR: [14.40, 27.50 ms]. 12M; n = 10, median = 13.5 ms IQR: [8.20, 16.35 ms]. 18M; n = 7, median = 13.00 ms IQR: [9.60, 15.80 ms]. Statistics were performed with a Kruskal-Wallis ANOVA followed by Dunn’s post-test)^r^. Notably, the difference in spike-delay between pre-hearing and young adult neurons was not significant. This behaviour therefore does not correlate with the known role of *I*_A_ in the generation of this delay (**Shibata et al., 2000**), and the progressive reduction of *I*_A_ during the onset of hearing cannot fully explain these spike-delay findings alone (**Figure 2F**)^b^. Comparatively, the firing rate of iLOC neurons over the course of the 250 ms stimulus increased with age when normalised to the current threshold (**Figure 5K**), which when compared at +50 pA revealed a significant increase during the onset of hearing and between young-adult and aged mice at 12 months. No statistical significance was observed between 12- and 18-month old ageing groups (*P* < 0.0001. <P12; n = 24, median = 28 Hz IQR: [20, 43 Hz]. 1-3M; n = 50, median = 48 Hz IQR: [40, 56 Hz]. 12M; n = 9, median = 72 Hz IQR: [60, 76 hZ]. 18M; n = 5, median = 68 Hz IQR: [56, 74 Hz]. Statistics were performed with a Kruskal-Wallis ANOVA followed by Dunn’s post-test)^s^.

**Figure 5.**
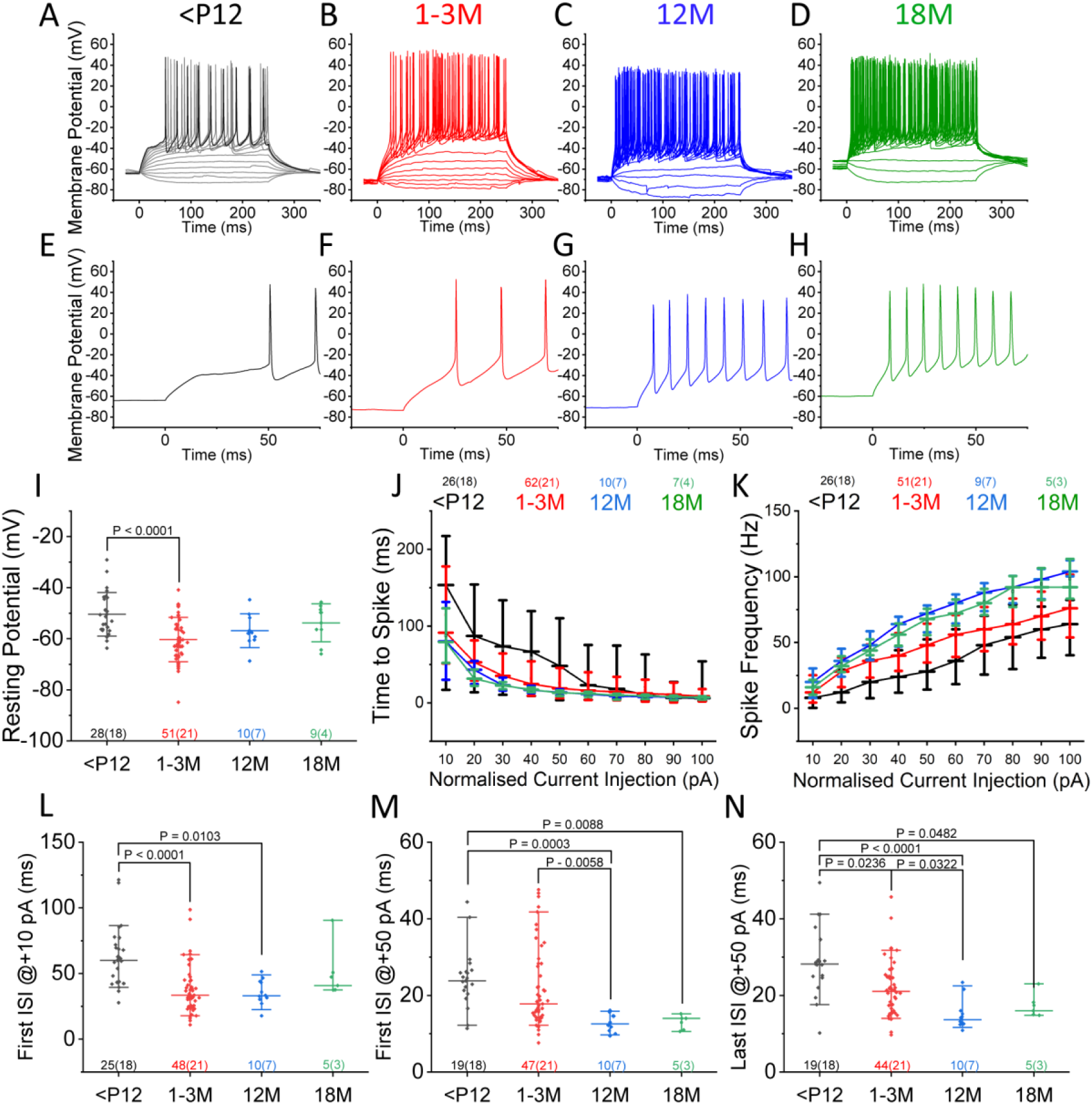
iLOC neuron firing rate increases and time to spike decreases with age. ***A-D***, Example voltage recordings of iLOC neurons from pre-hearing (***A*** *-* <P12; P6-P11), young adult (***B*** – 1-3M; P24-P89), 12 months (***C*** – 12M; P366-P416), and 18 months old (***D*** – 18M; P557-P564) mice in response to current injections from -20 pA to +100 pA in 10 pA steps. ***E-H***, Same example traces from ***A-D*** zoomed in to highlight the difference in delay between +100 pA current stimulus onset and the first action potential. ***I***, iLOC resting membrane potential calculated from the type of traces shown in Figure 4A-D prior to the onset of the current stimulus. ***J-K***, Time to the peak of the first action potential (***J***) and spiking frequency (***K***) for the four age groups where 10 pA normalised current injection is the rheobase for each neuron. ***L***, Time between the first two spikes of a +10 pA depolarising current injection, normalised such that 10 pA is the minimum current injection required to elicit at least two spikes. ***M***, First inter-spike interval in response to a +50 pA normalised current injection. ***N***, Time between the final two spikes of a 250 ms current injection step at +50 pA, normalised such that 10 pA is the minimum current injection required to elicit at least two spikes. Statistics were performed with a one-way ANOVA followed by Tuckey’s post-test or a Kruskal-Wallis ANOVA followed by Dunn’s post-test, with exact *P* values indicated over each comparison. The number of iLOC neurons and mice are shown beneath the data in the format; iLOC neurons (mice).

In response to weak current injection of +10 pA the first inter-spike interval (ISI) decreased during the onset of hearing, but not with ageing, correlating with the changes in *I*_A_ (**Figure 5L**)^t^. In response to stronger stimuli of +50 pA there was no difference between the first ISI of pre-hearing and young adult iLOC neurons, perhaps as the upregulated channel (Kv2) is too slow contribute to these ISIs at the very start of current injection (**Figure 5M**)^u^. However, with ageing and the upregulation of fast Kv3 channels, iLOC neurons showed a shorter first ISI relative younger neurons in response to larger current injection at +50 pA (**Figure 5M**)^u^. In contrast, the last ISI in response to stronger stimuli decreased with both the onset of hearing and ageing (**Figure 5N**)^v^. Taken together, iLOC action potential characteristics shift in a complicated manner with the onset of hearing and ageing, though trend towards an increased action potential output, showing a shorter spiking delay and a higher frequency for a given stimulus with no apparent reversion in the trend in ageing animals. Assuming iLOC neurons remain primarily cholinergic with age, this could represent an increased excitatory output to the cochlear relative to excitatory input to the iLOC neurons, perhaps facilitating increased gain with regards to activity of the spiral ganglion neurons.

Given the changes in the outward potassium currents and spiking characteristics, we then investigated whether the kinetics of individual spikes in iLOC neurons change with age. By injecting a depolarising current, action potentials from iLOC neurons were recorded near their threshold (**Figure 6A-D**, with corresponding phase plots shown in **6E-H**). With the onset of hearing and ageing, action potentials became faster (**Figure 6I**)^w^, which was not due to a faster rise time constant (**Figure 6J**)^x^, but due to a significantly reduced repolarisation time (**Figure 6K**)^y^. Despite the significant shift in *I*_Na_ size (**Figure 3H**)^f^ and peak voltage (**Figure 3I**)^g^, the spike voltage threshold, calculated from phase plots as shown in **Figure 6E**, did reveal an overall effect of age with a Kruskal-Wallis ANOVA but no significant comparisons were resolved with Dunn’s post-test (*P* = 0.0301. <P12; n = 26, median = -15.232 mV IQR: [-19.019, -7.460 mV]. 1-3M; n = 50, median = -16.327 mV IQR: [-19.772, -9.786 mV]. 12M; n = 10, median = -21.415 mV IQR: [-23.465, -16.574]. 18M; n = 8, median = -19.627 mV IQR: [-28.963, -11.564 mV])^z^. In contrast, action potential height showed no effect of age with the same statistical approach (*P* = 0.9781. <P12; n = 25, median = 75.546 mV IQR: [67.217, 81.139 mV]. 1-3M; n = 47, median = 75.498 mV IQR: [69.058, 79.332mV]. 12M; n = 9, median = 75.137 mV IQR: [65.418, 81.516 mV]. 18M; n = 7, median = 76.011 mV IQR: [65.762, 76.352 mV])^aa^. Input resistance calculated between the -82 and -72 voltage steps, as a measure of passive membrane properties, significantly increased after the onset of hearing, but did not change further with ageing and no other comparison was significant (**Figure 6L**)^ab^. In line with the literature, and despite the shift in input resistance, iLOC rheobase did not shift during post-natal development or in 12 month old animals, but did trend toward lower values with age, becoming significant when comparing pre-hearing iLOCs to those from 18 month old animals (**Figure 6M**)^ac^. Thus, in combination with the time to spike data, indicates that iLOC neurons become more excitable during post-natal development by requiring a larger charge to fire, but possess a stable rheobase until much older ages (**Figure 6M**)^ac^.

**Figure 6.**
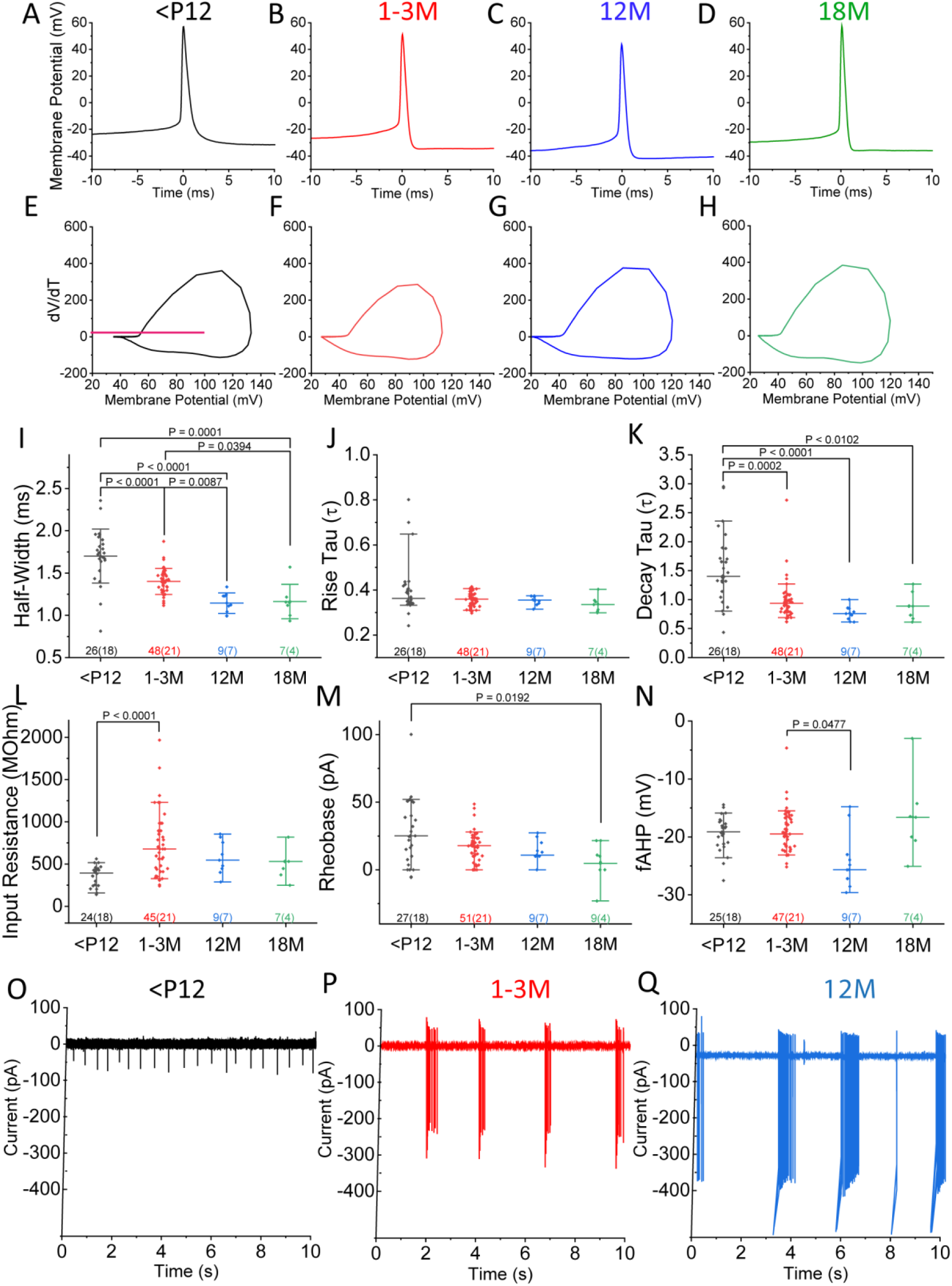
With age iLOC spikes have shorter half-widths due to faster repolarisation. ***A-D***, Example voltage recordings of iLOC neuron action potentials taken at the rheobase from pre-hearing (***A*** *-* <P12; P6-P11), young adult (***B*** – 1-3M; P24-P89), 12 months (***C*** – 12M; P366-P416), and 18 months old (***D*** – 18M; P557-P564) animals. ***E-H***, Example phase-plots of the action potential examples shown in ***A-D***, with the magenta line in ***E*** representing a dV/dT value of 20, from which the voltage threshold values were calculated from each cell. Action potential traces were used to calculate action potential half-widths (***I***), the time constant of the depolarisation rising phase (***J***), and the time constant of the falling repolarisation phase (***K***) utilising Clampfit software fitted as the sum of two exponentials and calculated 5% from the peak. ***L***, Input resistance values were calculated from the traces described in Figure 3 as the resistance between the -82 and -72 voltage steps, as a measure of passive membrane properties. Action potential recordings were taken at the rheobase of each neuron which was recorded (***M***), as was the minimal fast-after hyperpolarisation (fAHP) reached following the spike (***N***). ***O-Q***, Example cell-attached recordings of iLOC neurons in pre-hearing (***O,*** P11, n=2 cells from 1 animal), young adult (***P***, P32-P32, n=6 cells from 2 animals), and 12 months old mice (***Q***, P352-P404, n=3 cells from 2 animals) in the absence of current stimulation. All cell attached recordings were carried out with sucrose-based slicing solution. Statistics were performed with a one-way ANOVA followed by Tuckey’s post-test or a Kruskal-Wallis ANOVA followed by Dunn’s post-test, with exact *P* values indicated over each comparison. The number of iLOC neurons and mice are shown beneath the data in the format; iLOC neurons (mice).

Given the known role of Kv2 and Kv3 in shaping fast after-hyperpolarisation (fAHP), this was then investigated and measured as the lowest voltage reached immediately after an action potential. The fAHP was stable between pre-hearing and young-adult iLOC neurons, but became significantly more negative relative to the spiking voltage threshold at 12 months relative to young-adult animals (**Figure 6N**)^ad^. Notably, no statistical significance was found between the young-adult and 18-month old group. Altogether, the changes in spiking time-course align with the increased size of the overall outward potassium current (**Figure 2H-I**)^d^ as well as the known role of Kv2 within the superior olivary complex (**Tong et al., 2013**). Moreover, the more negative fAHP values in 12 month old ageing animals (**Figure 6N**)^ad^ is in agreement with the increase in rapid potassium (Kv3) current which is known to shape fAHP (**Figure 3F**)^e^ (**Rudy and McBain, 2001**), though this fast current alone cannot explain why fAHP does not remain significantly decreased at 18 months of age.

The absence of spontaneous activity in iLOC neurons in the whole-cell patch configuration (**Figure 6A-D**, **Figure 6M**)^ac^ was likely related to an artificial effect of the intra-cellular solution. As, when utilising cell-attached configurations and maintaining the neuronal intracellular environment, iLOC neurons fired spontaneously and demonstrated the shift in firing pattern reported by **Hong et al., (2022) (Figure 6O-Q**). We originally hypothesised that the firing pattern of aged iLOC neurons would be more similar to immature iLOCs (**Figure 6O**) than young adult neurons (**Figure 6P**) given that in aged mice, evidence suggests the iLOC synapses begin targeting the IHC somas similar to immature medial olivocochlear neurons in the pre-hearing cochlea. However, we found from limited pilot experiments that aged iLOCs appeared to retain bursting activity similar to 1-3M iLOC neurons (**Figure 6Q**), correlating with the voltage clamp findings, though greater N-numbers are required to permit a more detailed analysis. Taken together, despite ageing resulting in a return of immature-like direct axo-somatic efferent connections onto IHCs, global iLOC biophysics and spiking patterns show no apparent reversion to an immature configuration (**Lauer et al., 2012; Zachary and Fuchs, 2015; Corns et al., 2018; Jeng et al., 2021**).

### IHC efferent re-innervation does not fully recreate ageing biophysical properties in iLOC neurons

Ageing has been associated with the rewiring of cholinergic efferent fibres within the cochlea, resulting in the formation of inhibitory axo-somatic synapses onto mature IHCs (**Lauer et al., 2012; Zachary and Fuchs, 2015; Jeng et al., 2021**), a feature normally restricted to the pre-hearing stage (**Glowatzki and Fuchs, 2000; Katz et al., 2004; Marcotti et al., 2004**). A similar change in adult iLOC wiring has also been reported in a conditional knockout mouse (*Myo7a^-/-^Myo15-Cre^+/-^*), in this strain adult IHCs progressively lose their mechano-electrical transduction due to *Myo7a* downregulation, and this results in IHC efferent re-innervation (**Corns et al., 2018**). In these mice, efferent cholinergic synapses can be visualised on adult IHCs by the presence of the post-synaptic SK2 potassium channel (**Figure 7A-B**), which ensure efferent synapses onto IHCs are inhibitory in nature following the initial calcium entry through nicotinic ACh receptors (**Elgoyhen et al., 1994; Glowatzki and Fuchs, 2000**). Because these newly formed synaptic connections have been suggested to arise at least in part from the iLOC neurons (**Jeng et al., 2021; O’Connor et al., 2024**), we examined whether this early model of ageing-like cochlear re-wiring elicited age-related biophysical changes in iLOC neuronal somata. To test this hypothesis, we investigated the biophysical properties of iLOC neurons in control and *Myo7a^-/-^xMyo15-Cre^+/-^*mice at around 4 months of age, which is well after the appearance of robust efferent IHC re-innervation (**Figure 7A-B**). Current recordings using the -82 mV pre-conditioning protocol (**Figure 7C-D)**, indicated no statistically significant change in membrane capacitance (**Figure 7E**)^ae^, size of both *I*_A_ (**Figure 7F**)^af^ and *I*_Na_ (**Figure 7G**)^ag^, and the potential of the peak *I*_Na_ (**Figure 7H**)^ah^. However, with *I*_A_ inactivated (**Figure 7I-J**) whilst the slow potassium current remained unchanged (**Figure 7K**)^ai^, the fast outward potassium current was significantly elevated in mutant animals relative to controls (**Figure 7L**)^aj^. Thus, despite eliciting robust efferent re-innervation in the cochlea, the only a subset of ageing features were recapitulated by the conditional deletion of Myo7a in voltage clamp.

**Figure 7.**
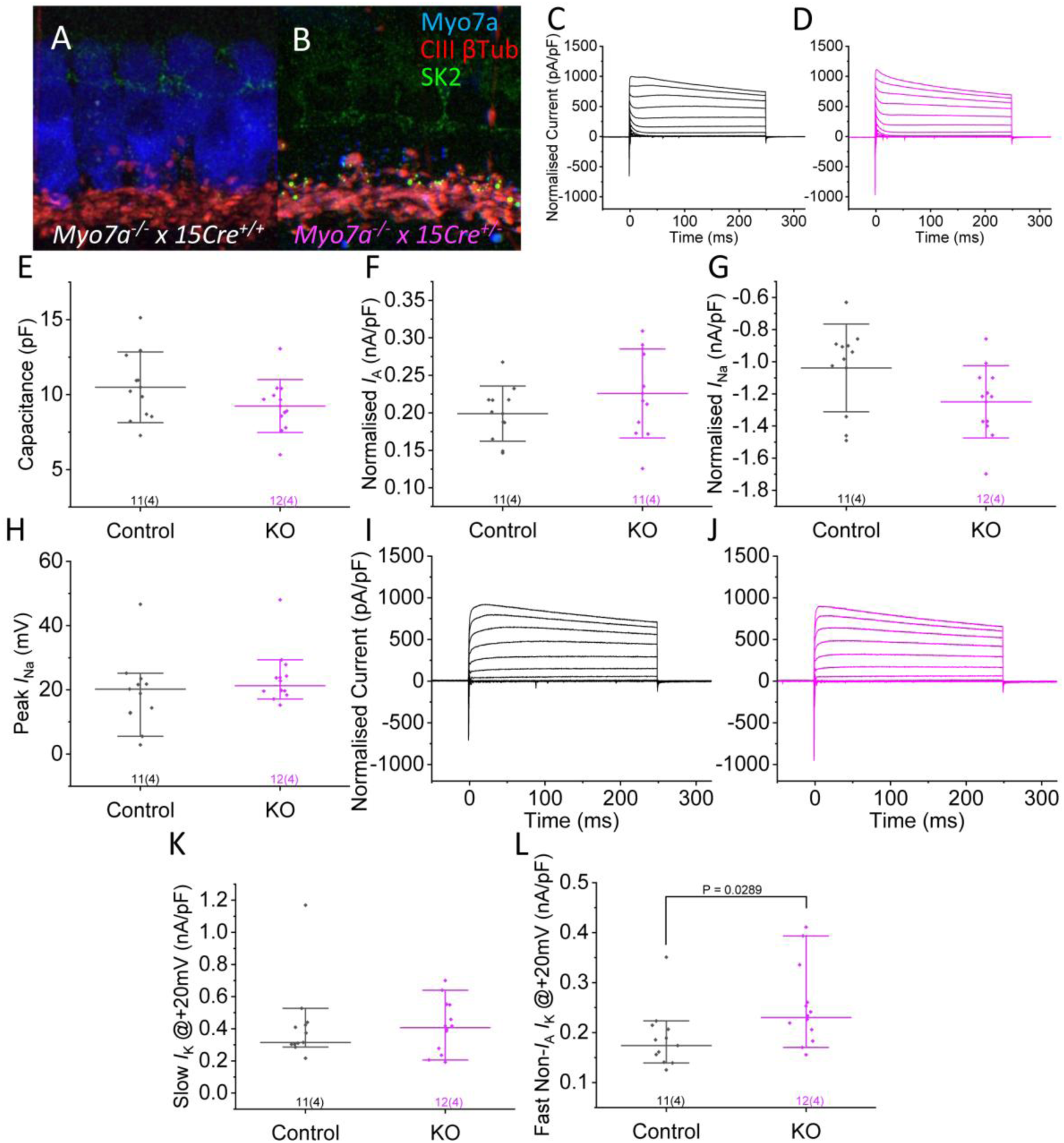
Post-natal hair cell specific deletion of *Myo7a* does not recapitulate immature or ageing iLOC voltage clamp biophysics. ***A-B***, Immunolabelling of P30 cochlear inner hair cells from either *Myo7a^-/-^ x Myosin 15-Cre^+/+^*control animals (***A***, n=3) or *Myo7a^-/-^ x Myosin 15-Cre^+/-^* hair cell specific knock-out (KO) mice (***B***, n=3). Inner hair cell re-innervation is evident as the re-appearance of SK2 puncta, and is already widely prevalent in P30 animals. ***C-D***, Average total current recordings from P96-P115 control (***C***, n=11 cells from 4 animals) and *Myo7a* conditional deletion (***D***, n=12 cells from 4 animals) iLOC neurons in response to voltage steps from -62 mV to +68 mV in 10 mV increments normalised to cell capacitance. Similar to the measurements made in Figure 2, iLOC capacitance (***E***), *I*_A_ density (***F***), density of *I*_Na_ (***G***), and the membrane potential at which *I*_Na_ is largest (***H***). ***I-J***, Average total current recordings from P106-P115 control (***I***) and deletion (***J***) iLOC neurons from -62 mV to +68mV in 10 mV increments normalised to cell capacitance. Size of the slow (***K***) and fast (***L***) non-*I*_A_ outward potassium current at +20 mV measured as in Figure 3. Statistics were performed with a Student’s t-test or Mann-Whitney U-test, with exact *P* values indicated over each comparison if statistically significant. The number of iLOC neurons and mice are shown beneath the data in the format; iLOC neurons (mice).

### IHC specific deletion of *Myo7a* leads to iLOC neurons in adult mice with limited immature spiking characteristics

To investigate if IHC dysfunction and re-innervation elicited ageing-like changes in iLOC spiking characteristics, we compared voltage responses from adult iLOC neurons from control and littermate *Myo7a* conditional KO animals. When current-clamp recordings were carried out with current injection from -20 to +100 pA iLOC neurons from both genotypes spiked readily (**Figure 8A-B**), with no statistically significant difference in input resistance (**Figure 8C**)^ak^ or voltage threshold (*P* = 0.5095. Control; n = 10, median = -18.909 mV IQR: [-35.632, -15.668 mV]. KO; n = 12, median = -19.952 mV IQR: [-37.718, -16.976 mV]. Statistics were performed with a Mann-Whitney U-test.)^al^. Spiking activity in response to depolarising current injection (**Figure 8A-B**) showed a significantly increased delay in the first ISI at lower stimulus intensities in *Myo7a* deficient mice (**Figure 8D**)^am^, reminiscent of the profile observed in pre-hearing iLOC neurons (**Figure 5L**)^t^. However, the ISIs in response to higher stimulus intensities of +50 pA were unaffected between genotypes at both the start (**Figure 8E**)^an^ and end of the current injection (**Figure 8F**)^ao^. Spike kinetics measured from analogue current injection similar to those shown in **Figure 6** indicated no significant difference between control and *Myo7a* conditional KO animals for either action potential half-width (**Figure 8G**)^ap^ or the decay timing constant (**Figure 8H**)^aq^. The resting membrane potential, however, significantly shifted to more depolarised values in *Myo7a* deficient mice, which together with the low-stimulus ISI (**Figure 8D**)^am^ may indicate a degree of reversion to a more immature firing profile in these adult iLOC neurons (**Figure 8I**)^ar^. This is, however, in confliction with the apparent up-regulation of the fast-potassium current seen in voltage-clamp configuration (**Figure 7J**)^aj^ that is associated with ageing-like biophysics (**Figure 3F**)^e^. Overall, iLOC neurons from *Myo7a* deficient animals fail to show either a full ageing or immature set of biophysics, relative to the pronounced shifts seen during the normal maturation and ageing life-course in 6N wild-type animals.

**Table 1-1.** Statistical Summary.

|  |  |  |
| --- | --- | --- |
| **<sup>a</sup>Figure 2E – Capacitance**<br>N = 122<br>4 Age groups | Normally Distributed.<br>One-Way ANOVA with | **Overall Main Effect:**<br>$F(3,118) = 29.48, P < 0.0001, R^2 = 0.43$<br>**Pairwise Comparisons:** |
|  | Tuckey's post-test. | • **<P12 vs 1-3M:** $q(118) = 11.57$, Adjusted $P < 0.0001$, 95% CI = [-8.27, -4.27]<br>• **12M vs <P12:** $q(118) = 20.23$, Adjusted $P < 0.0001$, 95% CI = [-9.80, -4.61]<br>• **12M vs 1-3M:** $q(118) = 1.44$, Adjusted $P = 0.7411$, 95% CI = [-3.32, 1.46]<br>• **18M vs <P12:** $q(118) = 9.59$, Adjusted $P < 0.0001$, 95% CI = [-9.68, -4.30]<br>• **18M vs 1-3M:** $q(118) = 1.07$, Adjusted $P = 0.8746$, 95% CI = [-3.21, 1.77]<br>• **18M vs 12M:** $q(118) = 0.26$, Adjusted $P = 0.9978$, 95% CI = [-2.78, 3.20] |
| <sup>b</sup>**Figure 2F – $I_A$ size**<br>N = 111<br>4 Age groups | Normally Distributed.<br>One-Way ANOVA with Tuckey's post-test. | **Overall Main Effect:**<br>$F(3,107) = 7.10$, $P = 0.0002$, $R^2 = 0.17$<br>**Pairwise Comparisons:**<br>• **<P12 vs 1-3M:** $q(107) = 5.81$, Adjusted $P = 0.0004$, 95% CI = [-0.12, -0.03]<br>• **12M vs <P12:** $q(107) = 5.41$, Adjusted $P = 0.0012$, 95% CI = [-0.15, -0.03]<br>• **12M vs 1-3M:** $q(107) = 1.17$, Adjusted $P = 0.8418$, 95% CI = [-0.07, 0.04]<br>• **18M vs <P12:** $q(107) = 3.34$, Adjusted $P = 0.0903$, 95% CI = [-0.12, 0.01]<br>• **18M vs 1-3M:** $q(107) = 1.00$, Adjusted $P = 0.8946$, 95% CI = [-0.04, 0.07]<br>• **18M vs 12M:** $q(107) = 1.74$, Adjusted $P = 0.6073$, 95% CI = [-0.04, 0.10] |
| <sup>c</sup>**Figure 2G – $I_A$ Half-inactivation**<br>N = 111<br>4 Age groups | Normally Distributed.<br>One-Way ANOVA with Tuckey's post-test. | **Overall Main Effect:**<br>$F(3,107) = 3.43$, $P = 0.0196$, $R^2 = 0.09$<br>**Pairwise Comparisons:**<br>• **<P12 vs 1-3M:** $q(107) = 1.72$, Adjusted $P = 0.6181$, 95% CI = [-2.40, 6.60] |
|  |  | • **12M vs <P12:** $q(107) = 0.63$, Adjusted $P = 0.9708$, 95% CI = [-5.00, 7.04]<br>• **12M vs 1-3M:** $q(107) = 0.72$, Adjusted $P = 0.9562$, 95% CI = [-6.57, 4.41]<br>• **18M vs <P12:** $q(107) = 2.85$, Adjusted $P = 0.1879$, 95% CI = [-10.89, 1.29]<br>• **18M vs 1-3M:** $q(107) = 4.49$, Adjusted $P = 0.0104$, 95% CI = [-12.48, -1.22]<br>• **18M vs 12M:** $q(107) = 3.09$, Adjusted $P = 0.1347$, 95% CI = [-12.67, 1.13] |
| <sup>d</sup>**Figure 2I – Sustained current at 20 mV**<br>N = 118<br>4 Age groups | Not Normally Distributed.<br>Kruskal-Wallis ANOVA with Dunn's post-test. | **Overall Main Effect:**<br>$\chi^2(3) = 46.69$, $P < 0.0001$, $\eta^2 = 0.37$<br>**Pairwise Comparisons:**<br>• **<P12 vs 1-3M:** $Z = -3.92$, Mean Rank Diff = -31.21, Adjusted $P = 0.0005$<br>• **<P12 vs 12M:** $Z = -6.46$, Mean Rank Diff = -65.65, Adjusted $P < 0.0001$<br>• **<P12 vs 18M:** $Z = -4.57$, Mean Rank Diff = -48.11, Adjusted $P = 0.0003$<br>• **1-3M vs 12M:** $Z = -3.77$, Mean Rank Diff = -34.44, Adjusted $P = 0.0010$<br>• **1-3M vs 18M:** $Z = -1.78$, Mean Rank Diff = -16.90, Adjusted $P = 0.4542$<br>• **12M vs 18M:** $Z = 1.54$, Mean Rank Diff = 17.54, Adjusted $P = 0.7468$ |
| <sup>e</sup>**Figure 3F – Fast $I_K$ current at 20 mV**<br>N = 122<br>4 Age groups | Normally Distributed.<br>One-Way ANOVA with Tukey's post-test. | **Overall Main Effect:**<br>$F(3,118) = 23.98$, $P < 0.0001$, $R^2 = 0.38$<br>**Pairwise Comparisons:**<br>• **<P12 vs 1-3M:** $q(118) = 1.66$, Adjusted $P = 0.6452$, 95% CI = [-0.07, 0.03]<br>• **12M vs <P12:** $q(118) = 8.35$, Adjusted $P < 0.0001$, 95% CI = [0.08, 0.21]<br>• **12M vs 1-3M:** $q(118) = 10.45$, Adjusted $P < 0.0001$, 95% CI = [0.11, 0.22] |
|  |  | • **18M vs <P12:** $q(118) = 5.74$, Adjusted $P = 0.0005$, 95% CI = [0.04, 0.17]<br>• **18M vs 1-3M:** $q(118) = 7.52$, Adjusted $P < 0.0001$, 95% CI = [0.06, 0.18]<br>• **18M vs 12M:** $q(118) = 2.08$, Adjusted $P = 0.4575$, 95% CI = [-0.11, 0.03] |
| <sup>f</sup>**Figure 3H – Sodium current size**<br>N = 122<br>4 Age groups | Not Normally Distributed.<br>Kruskal-Wallis ANOVA with Dunn's post-test. | **Overall Main Effect:**<br>$\chi^2(3) = 20.92$, $P = 0.0002$, $\eta^2 = 0.15$<br>**Pairwise Comparisons:**<br>• **<P12 vs 1-3M:** $Z = 2.83$, Mean Rank Diff = 22.33, Adjusted $P = 0.0279$<br>• **<P12 vs 12M:** $Z = 4.42$, Mean Rank Diff = 45.24, Adjusted $P < 0.0001$<br>• **<P12 vs 18M:** $Z = 1.08$, Mean Rank Diff = 11.50, Adjusted $P = 1.0000$<br>• **1-3M vs 12M:** $Z = 2.43$, Mean Rank Diff = 22.91, Adjusted $P = 0.0908$<br>• **1-3M vs 18M:** $Z = -1.10$, Mean Rank Diff = -10.83, Adjusted $P = 1.0000$<br>• **12M vs 18M:** $Z = -2.86$, Mean Rank Diff = -33.74, Adjusted $P = 0.0256$ |
| <sup>g</sup>**Figure 3I – Sodium current peak voltage**<br>N = 122<br>4 Age groups | Normally Distributed.<br>One-Way ANOVA with Tukey's post-test. | **Overall Main Effect:**<br>$F(3,118) = 7.98$, $P < 0.0001$, $R^2 = 0.17$<br>**Pairwise Comparisons:**<br>• **<P12 vs 1-3M:** $q(118) = 1.17$, Adjusted $P = 0.8406$, 95% CI = [-9.99, 5.17]<br>• **12M vs <P12:** $q(118) = 4.91$, Adjusted $P = 0.0039$, 95% CI = [-22.97, -3.28]<br>• **12M vs 1-3M:** $q(118) = 4.35$, Adjusted $P = 0.0136$, 95% CI = [-19.78, -1.65]<br>• **18M vs <P12:** $q(118) = 5.38$, Adjusted $P = 0.0013$, 95% CI = [-25.08, -4.68] |
|  |  | • **18M vs 1-3M:** $q(118) = 4.86$, Adjusted $P = 0.0045$, 95% CI = [-21.92, -3.02]<br>• **18M vs 12M:** $q(118) = 0.57$, Adjusted $P = 0.9777$, 95% CI = [-21.92, 9.59] |
| <sup>h</sup>**Inward current size**<br>N = 73<br>4 Age groups | Not Normally Distributed.<br>Kruskal-Wallis ANOVA with Dunn's post-test. | **Overall Main Effect:**<br>$\chi^2(3) = 6.30$, $P = 0.0977$, $\eta^2 = 0.05$<br>**Pairwise Comparisons:**<br>• **<P12 vs 1-3M:** $Z = -0.40$, Mean Rank Diff = -2.43, Adjusted $P = 1.0000$<br>• **<P12 vs 12M:** $Z = 1.92$, Mean Rank Diff = 16.67, Adjusted $P = 0.3260$<br>• **<P12 vs 18M:** $Z = 0.57$, Mean Rank Diff = 6.11, Adjusted $P = 1.0000$<br>• **1-3M vs 12M:** $Z = 2.44$, Mean Rank Diff = 19.09, Adjusted $P = 0.0870$<br>• **1-3M vs 18M:** $Z = 0.85$, Mean Rank Diff = 8.54, Adjusted $P = 1.0000$<br>• **12M vs 18M:** $Z = -0.89$, Mean Rank Diff = -10.56, Adjusted $P = 1.0000$ |
| <sup>i</sup>**Figure 3.1 - Relationship between voltage clamp metrics and tonotopy.** | Scatter plots against tonotopic axis.<br>Linear regression analysis with Bonferonni correction for multiple parameters. | **Linear Regression Parameter:**<br>• **Capacitance vs Position:** $N = 42$, $R^2 = -0.02$, $F(1, 40) = 0.00$, Adjusted $P = 1.0000$<br>• **Normalized $I_A$ vs Position:** $N = 41$, $R^2 = -0.01$, $F(1,39) = 0.47$, Adjusted $P = 1.0000$<br>• **$I_A$ Half-Inactivation vs Position:** $N = 39$, $R^2 = 0.11$, $F(1,39) = 6.00$, Adjusted $P = 0.1323$<br>• **Normalized $I_{Na}$ Size vs Position:** $N = 42$, $R^2 = -0.02$, $F(1,40) = 0.00$, Adjusted $P = 1.0000$<br>• **Peak $I_{Na}$ Voltage vs Position:** $N = 42$, $R^2 = -0.02$, $F(1,40) = 0.01$, Adjusted $P = 1.0000$<br>• **Slow $I_K$ Size vs Position:** $N = 42$, $R^2 = -0.02$, $F(1,40) = 0.32$, Adjusted $P = 1.0000$ |
|  |  | **Fast Non-$I_A$ $I_K$ Size vs Position:** $N = 42$, $R^2 = 0.00$, $F(1,40) = 1.00$, Adjusted $P = 1.0000$ |
| <sup>j</sup>**Figure 3.2D – Sucrose vs NMDG capacitance**<br>$N = 70$<br>3 Groups | Normally Distributed.<br>One-Way ANOVA with Tuckey's post-test. | **Overall Main Effect:**<br>$F(2,67) = 11.39$, $P < 0.0001$, $R^2 = 0.25$<br>**Pairwise Comparisons:**<br>**1M NMDG vs <P12:** $q(67) = 6.50$, Adjusted $P < 0.0001$, 95% CI = [-8.93, -2.81]<br>**1M Sucrose vs <P12:** $q(67) = 4.11$, Adjusted $P = 0.0136$, 95% CI = [-6.77, -0.65]<br>**1M Sucrose vs 1M NMDG:** $q(67) = 2.14$, Adjusted $P = 0.2923$, 95% CI = [-1.27, -5.59] |
| <sup>k</sup>**Figure 3.2E – Sucrose vs NMDG normalized $I_A$ size**<br>$N = 66$<br>3 Groups | Normally Distributed.<br>One-Way ANOVA with Tuckey's post-test. | **Overall Main Effect:**<br>$F(2,63) = 11.43$, $P < 0.0001$, $R^2 = 0.27$<br>**Pairwise Comparisons:**<br>**1M NMDG vs <P12:** $q(63) = 4.48$, Adjusted $P < 0.0066$, 95% CI = [-0.13, -0.02]<br>**1M Sucrose vs <P12:** $q(63) = 6.44$, Adjusted $P < 0.0001$, 95% CI = [-0.16, -0.05]<br>**1M Sucrose vs 1M NMDG:** $q(63) = 1.80$, Adjusted $P = 0.4165$, 95% CI = [-0.09, 0.03] |
| <sup>l</sup>**Figure 3.2F – Sucrose vs NMDG Sustained $I_K$ Size**<br>$N = 66$<br>3 Groups | Not Normally Distributed.<br>Kruskal-Wallis ANOVA with Dunn's post-test. | **Overall Main Effect:**<br>$\chi^2(2) = 17.74$, $P = 0.0001$, $\eta^2 = 0.25$<br>**Pairwise Comparisons:**<br>**<P12 vs 1M NMDG:** $Z = -3.74$, Mean Rank Diff = -21.35, Adjusted $P = 0.0005$<br>**<P12 vs 1M Sucrose:** $Z = -3.28$, Mean Rank Diff = 18.72, Adjusted $P = 0.0031$<br>**1M NMDG vs 1M Sucrose:** $Z = 0.42$, Mean Rank Diff = 2.63, Adjusted $P = 1.0000$ |
| <sup>m</sup>**Figure 3.2G – Sucrose vs NMDG Fast $I_K$ Current at 20 mV**<br>$N = 70$<br>3 Groups | Normally Distributed.<br>One-Way ANOVA with Tuckey's post-test. | **Overall Main Effect:**<br>$F(2,67) = 0.99$, $P = 0.3765$, $R^2 = 0.03$<br>**Pairwise Comparisons:**<br>**1M NMDG vs <P12:** $q(67) = 1.73$, Adjusted $P = 0.4438$, 95% CI = [-0.07, 0.02] |
|  |  | • **1M Sucrose vs <P12:** $q(67) = 1.56$, Adjusted $P = 0.5158$, 95% CI = [-0.07, -0.02]<br>• **1M Sucrose vs 1M NMDG:** $q(67) = 0.15$, Adjusted $P = 0.9936$, 95% CI = [-0.05, 0.05] |
| <sup>n</sup>**Figure 3.2H – Sucrose vs NMDG $I_{Na}$ Current Size**<br>N = 70<br>3 Groups | Not Normally Distributed.<br>Kruskal-Wallis ANOVA with Dunn's post-test. | **Overall Main Effect:**<br>$\chi^2(2) = 5.58$, $P = 0.0616$, $\eta^2 = 0.05$<br>**Pairwise Comparisons:**<br>• **<P12 vs 1M NMDG:** $Z = 1.64$, Mean Rank Diff = 9.68, Adjusted $P = 0.3018$<br>• **<P12 vs 1M Sucrose:** $Z = 2.19$, Mean Rank Diff = 12.89, Adjusted $P = 0.0863$<br>• **1M NMDG vs 1M Sucrose:** $Z = 0.49$, Mean Rank Diff = 3.21, Adjusted $P = 1.0000$ |
| <sup>o</sup>**Figure 4D – Fast Current Ratio with 1mM TEA**<br>N = 8<br>3 Groups | Normally Distributed paired values with missing values.<br>Paired Sample t-tests with Bonferonni correction for multiple comparisons | **Pairwise Drug Effects:**<br>• **aCSF vs 1mM TEA:** $t(7) = 3.89$, Adjusted $P = 0.0119$, 95% CI = [0.06, 0.24]<br>• **1mM TEA vs Wash:** $t(5) = -5.00$, Adjusted $P = 0.0082$, 95% CI = [-0.38, -0.12] |
| <sup>p</sup>**Figure 4H – Fast Current with 1μM RY785**<br>N = 6<br>2 Groups | Normally Distributed<br>Student's t-test with Welch correction. | **Pairwise Comparison:**<br>$t(5) = -6.79$, $P = 0.0011$, 95% CI = [-0.51, -0.23] |
| <sup>q</sup>**Figure 5I – Resting Membrane Potential**<br>N = 98<br>4 Age groups | Normally Distributed.<br>One-Way ANOVA with Tuckey's post-test. | **Overall Main Effect:**<br>$F(3,94) = 8.81$, $P < 0.0001$, $R^2 = 0.22$<br>**Pairwise Comparisons:**<br>• **<P12 vs 1-3M:** $q(94) = 7.12$, Adjusted $P < 0.0001$, 95% CI = [-15.04, -4.76]<br>• **12M vs <P12:** $q(94) = 2.96$, Adjusted $P = 0.1627$, 95% CI = [-14.50, 1.61] |
|  |  | • **12M vs 1-3M:** $q(94) = 1.69$, Adjusted $P = 0.6321$, 95% CI = [-4.11, -11.02]<br>• **18M vs <P12:** $q(94) = 1.49$, Adjusted $P = 0.7182$, 95% CI = [-11.76, 5.00]<br>• **18M vs 1-3M:** $q(94) = 3.02$, Adjusted $P = 0.1426$, 95% CI = [-1.38, 14.43]<br>• **18M vs 12M:** $q(118) = 0.85$, Adjusted $P = 0.8544$, 95% CI = [-6.98, 13.12] |
| **<sup>r</sup>Time to spike at +50 pA**<br>N = 103<br>4 Age groups | Not Normally Distributed.<br>Kruskal-Wallis ANOVA with Dunn's post-test. | **Overall Main Effect:**<br>$\chi^2(3) = 14.52$, $P = 0.0023$, $\eta^2 = 0.12$<br>**Pairwise Comparisons:**<br>• **<P12 vs 1-3M:** $Z = 1.59$, Mean Rank Diff = 11.39, Adjusted $P = 0.6765$<br>• **<P12 vs 12M:** $Z = 3.00$, Mean Rank Diff = 33.78, Adjusted $P = 0.0160$<br>• **<P12 vs 18M:** $Z = 2.97$, Mean Rank Diff = 38.16, Adjusted $P = 0.0177$<br>• **1-3M vs 12M:** $Z = 2.20$, Mean Rank Diff = 22.39, Adjusted $P = 0.1672$<br>• **1-3M vs 18M:** $Z = 2.25$, Mean Rank Diff = 26.77, Adjusted $P = 0.1478$<br>• **12M vs 18M:** $Z = 0.30$, Mean Rank Diff = 4.38, Adjusted $P = 1.0000$ |
| **<sup>s</sup>Figure 5K – Spike rate at 50 pA**<br>N = 88<br>4 Age groups | Not Normally Distributed.<br>Kruskal-Wallis ANOVA with Dunn's post-test. | **Overall Main Effect:**<br>$\chi^2(3) = 35.52$, $P < 0.0001$, $\eta^2 = 0.39$<br>**Pairwise Comparisons:**<br>• **<P12 vs 1-3M:** $Z = -3.77$, Mean Rank Diff = -23.79, Adjusted $P = 0.0010$<br>• **<P12 vs 12M:** $Z = -5.15$, Mean Rank Diff = -51.24, Adjusted $P < 0.0001$<br>• **<P12 vs 18M:** $Z = -3.89$, Mean Rank Diff = -48.62, Adjusted $P = 0.0006$<br>• **1-3M vs 12M:** $Z = -2.98$, Mean Rank Diff = -27.45, Adjusted $P = 0.0173$ |
|  |  | • **1-3M vs 18M:** $Z = -2.08$, Mean Rank Diff = -24.83, Adjusted $P = 0.2249$<br>• **12M vs 18M:** $Z = 0.18$, Mean Rank Diff = 2.62, Adjusted $P = 1.0000$ |
| <sup>t</sup>**Figure 5L – First inter-spike interval at +10 pA**<br>N = 88<br>4 Age groups | Not Normally Distributed.<br>Kruskal-Wallis ANOVA with Dunn's post-test. | **Overall Main Effect:**<br>$\chi^2(3) = 25.38$, $P < 0.0001$, $\eta^2 = 0.27$<br>**Pairwise Comparisons:**<br>• **<P12 vs 1-3M:** $Z = 4.83$, Mean Rank Diff = 30.46, Adjusted $P < 0.0001$<br>• **<P12 vs 12M:** $Z = 3.14$, Mean Rank Diff = 29.98, Adjusted $P = 0.0103$<br>• **<P12 vs 18M:** $Z = 0.93$, Mean Rank Diff = 11.58, Adjusted $P = 1.0000$<br>• **1-3M vs 12M:** $Z = -0.05$, Mean Rank Diff = -0.48, Adjusted $P = 1.0000$<br>• **1-3M vs 18M:** $Z = -1.57$, Mean Rank Diff = -18.88, Adjusted $P = 0.6946$<br>• **12M vs 18M:** $Z = -1.32$, Mean Rank Diff = -18.40, Adjusted $P = 1.0000$ |
| <sup>u</sup>**Figure 5M – First inter-spike interval at +50 pA**<br>N = 81<br>4 Age groups | Not Normally Distributed.<br>Kruskal-Wallis ANOVA with Dunn's post-test. | **Overall Main Effect:**<br>$\chi^2(3) = 22.48$, $P < 0.0001$, $\eta^2 = 0.25$<br>**Pairwise Comparisons:**<br>• **<P12 vs 1-3M:** $Z = 1.57$, Mean Rank Diff = 10.02, Adjusted $P = 0.7032$<br>• **<P12 vs 12M:** $Z = 4.03$, Mean Rank Diff = 37.06, Adjusted $P = 0.0003$<br>• **<P12 vs 18M:** $Z = 3.18$, Mean Rank Diff = 37.61, Adjusted $P = 0.0088$<br>• **1-3M vs 12M:** $Z = 3.30$, Mean Rank Diff = 27.04, Adjusted $P = 0.0058$<br>• **1-3M vs 18M:** $Z = 2.49$, Mean Rank Diff = 27.59, Adjusted $P = 0.0759$<br>• **12M vs 18M:** $Z = 0.04$, Mean Rank Diff = 0.55, Adjusted $P = 1.0000$ |
| <sup>v</sup>**Figure 5N – Last inter-spike interval at +50 pA**<br>N = 78<br>4 Age groups | Not Normally Distributed.<br>Kruskal-Wallis ANOVA with Dunn's post-test. | **Overall Main Effect:**<br>$\chi^2(3) = 22.79, P < 0.0001, \eta^2 = 0.27$<br>**Pairwise Comparisons:**<br>• **<P12 vs 1-3M:** $Z = 2.88$, Mean Rank Diff = 17.93, Adjusted $P = 0.0236$<br>• **<P12 vs 12M:** $Z = 4.52$, Mean Rank Diff = 40.03, Adjusted $P < 0.0001$<br>• **<P12 vs 18M:** $Z = 2.65$, Mean Rank Diff = 30.18, Adjusted $P = 0.0482$<br>• **1-3M vs 12M:** $Z = 2.78$, Mean Rank Diff = 22.10, Adjusted $P = 0.0322$<br>• **1-3M vs 18M:** $Z = 1.15$, Mean Rank Diff = 12.25, Adjusted $P = 1.0000$<br>• **12M vs 18M:** $Z = -0.79$, Mean Rank Diff = -9.85, Adjusted $P = 1.0000$ |
| <sup>w</sup>**Figure 6I – Action potential half-width**<br>N = 90<br>4 Age groups | Normally Distributed.<br>One-Way ANOVA with Tuckey's post-test. | **Overall Main Effect:**<br>$F(3,86) = 22.45, P < 0.0001, R^2 = 0.44$<br>**Pairwise Comparisons:**<br>• **<P12 vs 1-3M:** $q(86) = 8.04$, Adjusted $P < 0.0001$, 95% CI = [-0.44, -0.16]<br>• **12M vs <P12:** $q(86) = 9.38$, Adjusted $P < 0.0001$, 95% CI = [-0.78, -0.34]<br>• **12M vs 1-3M:** $q(86) = 4.60$, Adjusted $P = 0.0087$, 95% CI = [-0.46, -0.05]<br>• **18M vs <P12:** $q(86) = 8.24$, Adjusted $P < 0.0001$, 95% CI = [-0.78, -0.30]<br>• **18M vs 1-3M:** $q(86) = 3.84$, Adjusted $P = 0.0394$, 95% CI = [-0.47, -0.01]<br>• **18M vs 12M:** $q(86) = 0.24$, Adjusted $P = 0.9983$, 95% CI = [-0.27, 0.30] |
| <sup>x</sup>**Figure 6J – Action potential rise time constant**<br>N = 90<br>4 Age groups | Not Normally Distributed.<br>Kruskal-Wallis | **Overall Main Effect:**<br>$\chi^2(3) = 5.44, P = 0.1423, \eta^2 = 0.03$<br>**Pairwise Comparisons:** |
|  | ANOVA with Dunn's post-test. | • **<P12 vs 1-3M:** $Z = 1.24$, Mean Rank Diff = 7.90, Adjusted $P = 1.0000$<br>• **<P12 vs 12M:** $Z = 1.18$, Mean Rank Diff = 11.94, Adjusted $P = 1.000$<br>• **<P12 vs 18M:** $Z = 2.22$, Mean Rank Diff = 24.68, Adjusted $P = 0.16$<br>• **1-3M vs 12M:** $Z = 0.43$, Mean Rank Diff = 4.04, Adjusted $P = 1.0000$<br>• **1-3M vs 18M:** $Z = 1.59$, Mean Rank Diff = 16.78, Adjusted $P = 0.6735$<br>• **12M vs 18M:** $Z = 0.97$, Mean Rank Diff = 12.75, Adjusted $P = 1.0000$ |
| **<sup>y</sup>Figure 6K – Action potential decay time constant**<br>N = 90<br>4 Age groups | Not Normally Distributed.<br>Kruskal-Wallis ANOVA with Dunn's post-test. | **Overall Main Effect:**<br>$\chi^2(3) = 29.45$, $P < 0.0001$, $\eta^2 = 0.31$<br>**Pairwise Comparisons:**<br>• **<P12 vs 1-3M:** $Z = 4.19$, Mean Rank Diff = 26.64, Adjusted $P = 0.0002$<br>• **<P12 vs 12M:** $Z = 4.59$, Mean Rank Diff = 46.34, Adjusted $P < 0.0001$<br>• **<P12 vs 18M:** $Z = 3.14$, Mean Rank Diff = 34.91, Adjusted $P = 0.0102$<br>• **1-3M vs 12M:** $Z = 2.08$, Mean Rank Diff = 19.69, Adjusted $P = 0.2277$<br>• **1-3M vs 18M:** $Z = 0.78$, Mean Rank Diff = 8.27, Adjusted $P = 1.0000$<br>• **12M vs 18M:** $Z = -0.87$, Mean Rank Diff = -11.42, Adjusted $P = 1.0000$ |
| **<sup>z</sup>Voltage threshold**<br>N = 94<br>4 Age groups | Not Normally Distributed.<br>Kruskal-Wallis ANOVA with Dunn's post-test. | **Overall Main Effect:**<br>$\chi^2(3) = 8.94$, $P = 0.0301$, $\eta^2 = 0.07$<br>**Pairwise Comparisons:**<br>• **<P12 vs 1-3M:** $Z = 0.85$, Mean Rank Diff = 5.60, Adjusted $P = 1.0000$<br>• **<P12 vs 12M:** $Z = 2.57$, Mean Rank Diff = 26.08, Adjusted $P = 0.0612$ |
|  |  | • **<P12 vs 18M:** $Z = 1.95$, Mean Rank Diff = 21.45, Adjusted $P = 0.3106$<br>• **1-3M vs 12M:** $Z = 2.17$, Mean Rank Diff = 20.48, Adjusted $P = 0.1812$<br>• **1-3M vs 18M:** $Z = 1.53$, Mean Rank Diff = 15.86, Adjusted $P = 0.7614$<br>• **12M vs 18M:** $Z = -0.36$, Mean Rank Diff = -4.63, Adjusted $P = 1.0000$ |
| <sup>aa</sup>**Action potential spike height**<br>N = 88<br>4 Age groups | Not Normally Distributed.<br>Kruskal-Wallis ANOVA with Dunn's post-test. | **Overall Main Effect:**<br>$\chi^2(3) = 0.20$, $P = 0.9781$, $\eta^2 = -0.03$<br>**Pairwise Comparisons:**<br>• **<P12 vs 1-3M:** $Z = 0.24$, Mean Rank Diff = 1.51, Adjusted $P = 1.0000$<br>• **<P12 vs 12M:** $Z = 0.40$, Mean Rank Diff = 3.96, Adjusted $P = 1.0000$<br>• **<P12 vs 18M:** $Z = 0.28$, Mean Rank Diff = 3.10, Adjusted $P = 1.0000$<br>• **1-3M vs 12M:** $Z = 0.26$, Mean Rank Diff = 2.45, Adjusted $P = 1.0000$<br>• **1-3M vs 18M:** $Z = 0.15$, Mean Rank Diff = 1.59, Adjusted $P = 1.0000$<br>• **12M vs 18M:** $Z = -0.07$, Mean Rank Diff = -0.86, Adjusted $P = 1.0000$ |
| <sup>ab</sup>**Figure 6L – Input resistance**<br>N = 85<br>4 Age groups | Not Normally Distributed.<br>Kruskal-Wallis ANOVA with Dunn's post-test. | **Overall Main Effect:**<br>$\chi^2(3) = 24.39$, $P < 0.0001$, $\eta^2 = 0.26$<br>**Pairwise Comparisons:**<br>• **<P12 vs 1-3M:** $Z = -4.90$, Mean Rank Diff = -30.57, Adjusted $P < 0.0001$<br>• **<P12 vs 12M:** $Z = -2.35$, Mean Rank Diff = -22.67, Adjusted $P = 0.1126$<br>• **<P12 vs 18M:** $Z = -1.43$, Mean Rank Diff = -15.19, Adjusted $P = 0.9108$ |
|  |  | • **1-3M vs 12M:** $Z = 0.88$, Mean Rank Diff = 7.90, Adjusted $P = 1.0000$<br>• **1-3M vs 18M:** $Z = 1.53$, Mean Rank Diff = 15.38, Adjusted $P = 0.7505$<br>• **12M vs 18M:** $Z = 0.60$, Mean Rank Diff = 7.48, Adjusted $P = 1.0000$ |
| <sup>ac</sup>**Figure 6M – Rheobase**<br>N = 96<br>4 Age groups | Not Normally Distributed.<br>Kruskal-Wallis ANOVA with Dunn's post-test. | **Overall Main Effect:**<br>$\chi^2(3) = 9.57$, $P = 0.0226$, $\eta^2 = 0.07$<br>**Pairwise Comparisons:**<br>• **<P12 vs 1-3M:** $Z = 1.72$, Mean Rank Diff = 11.41, Adjusted $P = 0.5088$<br>• **<P12 vs 12M:** $Z = 1.60$, Mean Rank Diff = 17.13, Adjusted $P = 0.6585$<br>• **<P12 vs 18M:** $Z = 2.95$, Mean Rank Diff = 31.57, Adjusted $P = 0.0192$<br>• **1-3M vs 12M:** $Z = 0.57$, Mean Rank Diff = 5.72, Adjusted $P = 1.0000$<br>• **1-3M vs 18M:** $Z = 2.00$, Mean Rank Diff = 20.16, Adjusted $P = 0.2705$<br>• **12M vs 18M:** $Z = 1.10$, Mean Rank Diff = 14.44, Adjusted $P = 1.0000$ |
| <sup>ad</sup>**Figure 6N – fAHP**<br>N = 88<br>4 Age groups | Not Normally Distributed.<br>Kruskal-Wallis ANOVA with Dunn's post-test. | **Overall Main Effect:**<br>$\chi^2(3) = 8.73$, $P = 0.0332$, $\eta^2 = 0.07$<br>**Pairwise Comparisons:**<br>• **<P12 vs 1-3M:** $Z = -0.07$, Mean Rank Diff = -0.45, Adjusted $P = 1.0000$<br>• **<P12 vs 12M:** $Z = 2.44$, Mean Rank Diff = 24.22, Adjusted $P = 0.0884$<br>• **<P12 vs 18M:** $Z = -0.85$, Mean Rank Diff = -9.29, Adjusted $P = 1.0000$<br>• **1-3M vs 12M:** $Z = 2.65$, Mean Rank Diff = 24.67, Adjusted $P = 0.0477$<br>• **1-3M vs 18M:** $Z = -0.85$, Mean Rank Diff = -8.84, Adjusted $P = 1.0000$ |
|  |  | • **12M vs 18M:** $Z = -2.60$, Mean Rank Diff = -33.51, Adjusted $P = 0.0555$ |
| <sup>ae</sup>**Figure 7E – Control vs Myo7a capacitance**<br>N = 23<br>2 Groups | Normally Distributed.<br>Student's two-sample t-test with Welch correction. | **Pairwise Comparison:**<br>$t(18.49) = 1.44$, $P = 0.1675$, 95% CI = [-0.54, 3.05] |
| <sup>af</sup>**Figure 7F – Control vs Myo7a $I_A$ size**<br>N = 22<br>2 Groups | Normally Distributed.<br>Student's two-sample t-test with Welch correction. | **Pairwise Comparison:**<br>$t(16.71) = -1.28$, $P = 0.2191$, 95% CI = [-0.07, 0.02] |
| <sup>ag</sup>**Figure 7G – Control vs Myo7a $I_{Na}$ size**<br>N = 23<br>2 Groups | Normally Distributed.<br>Student's two-sample t-test with Welch correction. | **Pairwise Comparison:**<br>$t(19.48) = 2.03$, $P = 0.0581$, 95% CI = [-0.01, 0.43] |
| <sup>ah</sup>**Figure 7H – Control vs Myo7a Sodium current peak voltage**<br>N = 23<br>2 Groups | Not Normally Distributed.<br>Mann-Whitney U-test | **Pairwise Comparison:**<br>$U = 45$, $Z = -1.26$, $P = 0.2071$, $r = 0.26$ |
| <sup>ai</sup>**Figure 7K – Control vs Myo7a sustained current at +20 mV**<br>N = 23<br>2 Groups | Not Normally Distributed.<br>Mann-Whitney U-test | **Pairwise Comparison:**<br>$U = 59$, $Z = -0.40$, $P = 0.6891$, $r = 0.08$ |
| <sup>aj</sup>**Figure 7L – Control vs Myo7a fast current at +20 mV**<br>N = 23<br>2 Groups | Not Normally Distributed.<br>Mann-Whitney U-test | **Pairwise Comparison:**<br>$U = 30$, $Z = -2.18$, $P = 0.0289$, $r = 0.45$ |
| <sup>ak</sup>**Figure 8C – Control vs Myo7a input resistance**<br>N = 21<br>2 Groups | Not Normally Distributed.<br>Mann-Whitney U-test | **Pairwise Comparison:**<br>$U = 61.5$, $Z = 0.42$, $P = 0.6723$, $r = 0.09$ |
| <sup>al</sup>**Control vs Myo7a voltage threshold** | Not Normally Distributed. | **Pairwise Comparison:**<br>$U = 70.5$, $Z = 0.66$, $P = 0.5095$, $r = 0.14$ |
| N = 22 | Mann-Whitney U-test |  |
| <sup>am</sup>**Figure 8D – Control vs Myo7a first inter-spike interval at +10 pA**<br>N = 21<br>2 Groups | Normally Distributed.<br>Student's two-sample t-test with Welch correction. | **Pairwise Comparison:**<br>$t(16.45) = -3.19, P = 0.0056$ 95% CI = [-44.70, -7.19] |
| <sup>an</sup>**Figure 8E – Control vs Myo7a first inter-spike interval at +10 pA**<br>N = 21<br>2 Groups | Normally Distributed.<br>Student's two-sample t-test with Welch correction. | **Pairwise Comparison:**<br>$t(13.35) = -0.58, P = 0.5728$ 95% CI = [-4.77, 2.61] |
| <sup>ao</sup>**Figure 8F – Control vs Myo7a first inter-spike interval at +10 pA**<br>N = 20<br>2 Groups | Normally Distributed.<br>Student's two-sample t-test with Welch correction. | **Pairwise Comparison:**<br>$t(17.05) = -0.03, P = 0.9742$ 95% CI = [-6.36, 6.16] |
| <sup>ap</sup>**Figure 8G – Control vs Myo7a action potential half-width**<br>N = 22<br>2 Groups | Not Normally Distributed.<br>Mann-Whitney U-test | **Pairwise Comparison:**<br>$U = 71, Z = 0.69, P = 0.4887, r = 0.15$ |
| <sup>aq</sup>**Figure 8H – Control vs Myo7a decay time constant**<br>N = 22<br>2 Groups | Normally Distributed.<br>Student's two-sample t-test with Welch correction. | **Pairwise Comparison:**<br>$t(17.18) = 0.29, P = 0.7728$ 95% CI = [-0.12, 0.17] |
| <sup>ar</sup>**Figure 8I – Control vs Myo7a resting membrane potential**<br>N = 22<br>2 Groups | Normally Distributed.<br>Student's two-sample t-test with Welch correction. | **Pairwise Comparison:**<br>$t(19.91) = -3.01, P = 0.0069$ 95% CI = [-12.62, -2.15] |

**Figure 8.**
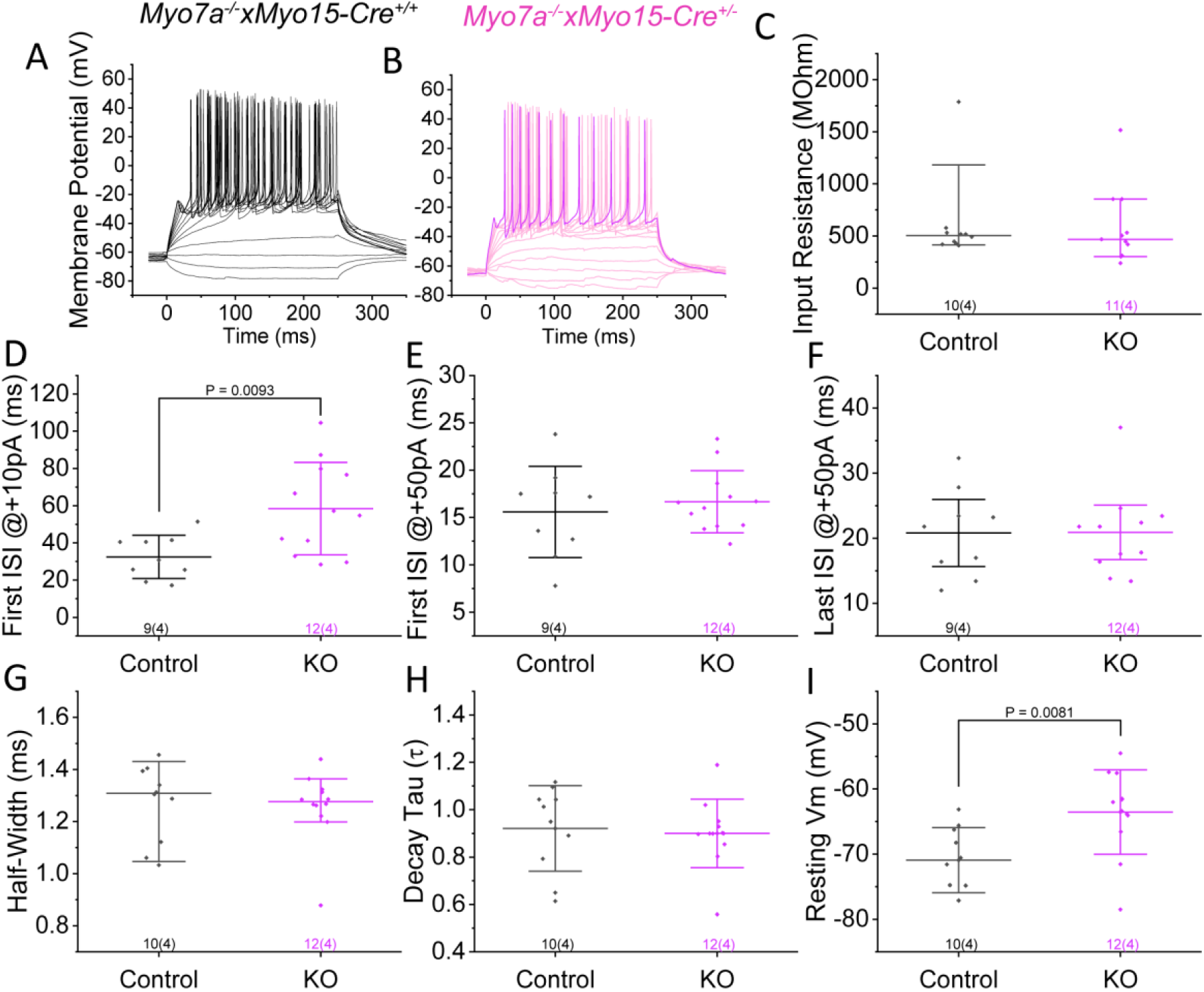
Post-natal hair cell specific deletion of *Myo7a* results in limited recapitulation of immature iLOC neurons in current clamp configuration. ***A-B***, Example voltage recordings from P96-P115 *Myo7a^-/-^ x Myosin 15-Cre^+/+^* control animals (***A***, n=9 cells from 4 animals) or *Myo7a^-/-^ x Myosin 15-Cre^+/-^* hair cell specific knock-out (KO) mice (***B***, n=12 cells from 4 animals) in response to current injection from -20 pA to +100 pA in increments of 10 pA. ***C***, Input resistance values calculated from the traces described in Figure 7, calculated as the resistance between the -82 and -72 voltage step. ***D-E***, First inter-spike interval after the onset of depolarising current injection at +10 pA (***D***) and +50 pA (***E***), normalised to the rheobase such that +10 pA is the lowest current injection that elicits two spikes. ***F***, Time between the last two spikes of an iLOC neuron following current injection at +50 pA normalised to the rheobase where +10 pA is the minimum current injection required to elicit at least two spikes. ***G-H***, iLOC action potential half-width (***G***) and decay time constant (***H***) values for the first action potential generated at the current injection threshold. ***I***, The resting membrane potential of iLOC neurons in the absence of current injection, averaged from the first 20 ms of traces such as ***A*** and ***B*** prior to the onset of the stimulus. Statistics were performed with a Student’s t-test or Mann-Whitney U-test, with exact *P* values indicated over each comparison. The number of iLOC neurons and mice are shown beneath the data in the format; iLOC neurons (mice).

## Discussion

Feedback to the cochlea from the brainstem remains poorly understood due to the severance of efferent fibres that occurs when the cochlea is dissected for *ex vivo* experimentation. As such, mechanistic insights into the functional characteristics of the efferent system can be achieved by investigating the neuronal somata within their resident nuclei. Here we have shown that iLOC neurons, through changes in Kv2, Kv3, Kv4 and Na_v_ mediated currents, adjust their excitability to generate and shape spontaneous bursting activity throughout life. Overall, these findings indicate that the biophysics of iLOC neurons progressively adapt with age allowing for a reduced action potential delay and a greater number of spikes for a given stimulus. In ageing mice, this enhanced excitatory output to SGNs may represent a compensatory mechanism, as iLOC neurons have been implicated in compensating for noise induced damage and enhance their bursting activity following noise insult (**Hong and Trussell, 2025; Sitko et al., 2025**). As such, during ageing iLOC neurons may provide progressively more excitatory cholinergic input to SGNs in order to maintain cochlear sensitivity. As the total activity of iLOC is driven by both intrinsic calcium-dependent bursting (**Hong et al., 2022**) and sound elicited input via the cochlear nucleus (**Doucet and Ryugo, 2003; Sterenborg et al., 2010; Romero and Trussell, 2021**), this could result in enhanced input to SGNs at rest or during sound stimulation.

### iLOC neuron biophysics change in a distinct manner as the hearing system matures and ages

One of the potassium current characteristic of iLOC neurons is the rapid inactivating A-type current (*I*_A_), which is carried by Kv4 subunits (**Birnbaum et al., 2004; Leijon and Magnusson, 2014**). *I*_A_ acts as a sub-threshold “brake” on action potentials and is responsible for the delay to the first action potential seen in iLOC neurons (**Hoffman et al., 1997; Sterenborg et al., 2010**). We found that the size of *I*_A_ in iLOC neurons is reduced during the onset of hearing, which resulted in a decrease of the first inter-spike interval, thus neurons became more readily excitable. iLOC spontaneous firing is calcium-dependent, and has been suggested to arise from prolonged activation of Cav1.3 calcium channels **(Hong and Trussell, 2025**). These channels activate at around the resting membrane potential, and push the neuron towards the voltage threshold until a spike is initiated.

This increased excitability was also due to the increased Kv2 mediated outward potassium current, which by shortening action potentials through faster repolarisation underpinned the progressive increase in action potential frequency. These results align with previous work in neurons of the medial and ventral nucleus of the trapezoid body (MNTB and VNTB respectively), where the knockdown of Kv2.2 channels resulted in reduced firing frequency (**Tong et al., 2013**). The VNTB contains the medial olivocochlear (MOC) efferent neurons, which suppress cochlear amplification via direct contacts onto cochlear outer hair cells (**Guinan, 2006**). It was found that disrupting Kv2.2 resulted in increased vulnerability to noise insult (**Tong et al., 2013**), and this was attributed solely to deficient MOC feedback. However, the work shown here indicates that post-hearing iLOC neurons also possess Kv2 channels (**Figure 2**,**6**). Thus, iLOC impairment may have contributed to this documented noise vulnerability, as iLOC neurons assist in threshold recovery following noise insult (**Sitko et al., 2025**). Kv2 upregulation is one of the major biophysical changes occurring in iLOC neurons during post-natal development (**Figure 2**), and the knock-down of Kv2 may impair normal function and revert iLOCs towards a more immature firing pattern. This may result in iLOCs that are unable to initiate bursts due to a more depolarised baseline and only firing once or twice per depolarisation (**Figure 6O-Q)**, though this interpretation requires experimental confirmation. Importantly, pharmacological blockade of Kv2 in immature iLOC neurons is required, to determine whether Kv2 is present at lower levels in pre-hearing animals or another slow-kinetics channel is operating, or whether slow opening Kv2-like channels are present only in adult neurons.

Alongside slow-activating Kv2, another major component of the outward potassium current in mature iLOC neurons is carried by Kv3 channels. This presence of Kv3 channels within iLOC neurons is unsurprising, as Kv3.4 was shown to be expressed in iLOC neuronal projections within the cochlea with immunolabelling (**Kim et al., 2020**). Whilst the current size of Kv3 did not change with the onset of hearing, ageing iLOC neurons appeared to strongly upregulate these rapid-activating Kv3 channels, further reducing action potential widths and increasing iLOC firing frequency for a given stimulus. Faster action potentials result in less net sodium channel inactivation per spike, possibly causing the longer spontaneous spike trains seen in aged iLOC neurons, though more cell-attached experiments are required for this to be robustly assessed (**Figure 6O-Q**; **Akemann and Knöpfel, 2006**).

Finally, we have shown here how the sodium current increased in size and peaked at lower voltages with age. Due to spike height and rise time being more related to channel kinetics than current size, we did not see any change in these values (**Hodgkin and Huxley, 1990**). However, an increase in sodium current density is known to be critical for maintaining spike capability during sustained action potential trains; making the neuron more resilient to cumulative sodium channel inactivation and still able to fire during longer bursts (**Raman and Bean, 1997; Khaliq et al., 2003**). As such, the increasing sodium current evident in iLOCs with post-natal development may be essential for the establishment and extension of spontaneous bursts.

Due to the decrease in cell surface area of iLOC neurons during the onset of hearing, an important consideration is whether the differences in biophysics seen in this work between pre-and post-hearing iLOC neurons are a result of changes in membrane ratio between the dendritic tree and soma. Kv4 channels that carry *I*_A_ (**Birnbaum et al., 2004**) reside primarily in the distal dendrites of neurons (**Hoffman et al., 1997**), whereas Kv2 and Kv3 channels reside proximal to the soma (**Trimmer, 2015**). If membrane area of the dendrites decreases relative to the soma between pre- and post-hearing stages, we would expect the Kv4 mediated current to decrease relative to Kv2 and Kv3. However, as Kv2 and Kv3 appear to be upregulated at distinct ages to one another (**Figure 2, 3**), this membrane area model cannot explain all the changes in biophysical profile we see here during post-natal development.

### Variations in the biophysical properties of iLOC neuron subpopulations

Here, iLOC neurons and their biophysical characteristics have been considered as a single population. However, recent findings utilising bulk RNAseq have identified two sub-groups, named LOC1 and LOC2 **(Frank et al., 2023**). Furthermore, some iLOC neurons upregulate tyrosine hydroxylase (TH) in response to noise exposure (**Wu et al., 2020**). As such, future work will be aimed at defining whether these different populations of iLOC neurons have distinct biophysical properties. Notably, LOC1 and LOC2 neurons tend to reside at opposite ends of the LSO (**Frank et al., 2023**), and as such any large changes in their biophysics would have likely been detected from our recordings (**Figure 4**).

On top of expression based sub-types, iLOC neurons may also be subdivided based on their synaptic targets in the cochlea, with this being the conventional type-I afferent terminals or, in aged mice, the additional but smaller number of direct axon-IHC contacts. As such, as iLOC neurons re-innervating IHCs directly likely makes up a small subset if iLOC neurons are indeed the re-innervating neuron, it is not possible to determine whether patched neurons were re-innervating an IHC.

### The mechanism by which the LSO responds to functional changes in the cochlea

In order to determine if age-related cochlear IHC efferent re-innervation was a driver for the shift in biophysics seen in ageing, we investigated iLOC neurons in animals with the conditional deletion of *Myo7a* from cochlear hair cells. Assuming re-innervation was the driver for ageing biophysics, and iLOC neurons are indeed the re-innervating neurons instead of medial olivocochlear neurons, we would expect the total outward potassium current, the fast outward potassium current, and the sodium current to be similarly up-regulated in the *Myo7a* model. Despite showing robust efferent re-innervation by a month of age, iLOC neurons from three months old *Myo7a x Myo15Cre* mice showed few biophysical shifts similar to those of ageing neurons, capturing only the increased fast potassium current. Current clamp also indicated no shared features with ageing iLOC neurons, and instead only some changes that are more consistent with an immature biophysical profile. This is perhaps unsurprising; as recent findings have indicated that when *Myo7a* is reintroduced via AAV gene therapy in *Myo7a* conditional knock-out animals, direct axo-somatic efferent contacts onto IHCs detach from only those cells expressing functional *Myo7a* (**O’Connor et al., 2024**). This suggests that IHC re-innervation itself is under local control within the cochlea, rather than a change that is occurring via an anterograde effect from the LSO. Given that it does not appear as though re-innervation cannot fully recapitulate the biophysical changes within the LSO, this suggests that iLOC neuron biophysics are responding to either a drop in afferent fibre activity induced by age-related changes within the hair cells or SGN fibre degeneration rather than hair cell transduction. Further studies are required to determine exactly what links cochlear activity or state with the feedback from the brainstem, and better understand the functional consequences on hearing.

Overall, it is important to note that whilst this study has provided novel insights into how iLOC neurons regulate their excitability within the LSO, *in vivo* methodologies are ultimately required to provide a clear picture as to exactly how this translates into synaptic activity onto the SGNs.

## Author contribution

The conceptualization, data collection, and analysis of this work were all carried out by AJC, who is also accountable for the integrity of the presented data and ensuring any questions related to data integrity are investigated and resolved adequately.

## Data availability statement

All data presented in this publication is available from the author upon reasonable request.

## Acknowledgements

The author thanks Mary Sherwood, Michelle Bird and Carol Pentland at the University of Sheffield animal facility for their assistance with mouse husbandry and care. Thanks also go to Niovi Voulgari, Catherine Gennery, and Matthew Hool for their assistance with genotyping. Special thanks go to Jing-Yi Jeng, Katherine Wood, and Walter Marcotti at the University of Sheffield for their advice and comments on the manuscript.

